# Individual Bird Identification by Modeling Temporal Structure in Bioacoustic Embeddings

**DOI:** 10.64898/2026.05.21.727031

**Authors:** Jonathan Gallego, Juan D. Martínez, José D. López

**Author notes:** Corresponding author *Email address:* (Jonathan Gallego).

## Abstract

1. Identifying individual animals from vocalizations is an emerging research area in computational bioacoustics. This non-invasive approach reduces reliance on physical capture and tagging for wildlife monitoring. Recent advances in this area leverage deep learning and bioacoustic foundation models adapted from species-level classifiers. However, these models typically rely on fixed-window inputs and may not fully capture temporal structure across extended or complex songs, which can contain information relevant to individual discrimination.
2. Here, we evaluate whether modeling sequences of pretrained bioacoustic embeddings improves acoustic individual identification. We developed a framework that integrates transfer-learned spectrotemporal representations from BirdNET with a lightweight long short-term memory network. Unlike static baselines that use either the first embedding or an average over all embeddings, our approach processes time-ordered embedding sequences, allowing the classifier to use information distributed across multiple windows.
3. We evaluated the system using 87,865 vocalizations from 352 individuals across seven species. We used four publicly available vocalization datasets with individual-level labels, covering durations from 0.76 s (short calls) to 27.6 s (prolonged songs). Across five random seeds with stratified partitions, the framework achieved mean test accuracies between 93.9% and 98.3% and macro-F1 scores ranging from 93.2% to 98.3%, without data augmentation. The clearest gains over the strongest static baseline were observed for the great tit and the great spotted kiwi, reaching +1.3 and +2.8 percentage points, respectively.
4. Our results indicate that the contribution of temporal sequence modeling depends on vocalization structure, rather than providing uniform evidence that chronological order drives performance. The benefit of recurrent aggregation was limited or absent for short calls but evident for long vocalizations with multiple informative windows. By combining bioacoustic foundation models with lightweight recurrent modeling, this approach provides a scalable, CPU-efficient tool for autonomous wildlife monitoring, particularly for species with extended or structurally complex vocalizations.

## 1. INTRODUCTION

Identifying individual animals from their vocalizations is an open and still-evolving frontier in bioacoustics. Although species-level identification is essential for biodiversity monitoring, acoustic data can also provide information about individual-level variation, behavior, and population processes (Knight et al., 2024). In particular, many bird species produce vocalizations that exhibit uniquely identifiable acoustic features, known as acoustic signatures (Budka et al., 2015; Terry et al., 2005), which can be leveraged to non-invasively monitor individuals, avoiding drawbacks associated with traditional marking techniques (Wilson and McMahon, 2006; Barron et al., 2010; Camacho et al., 2017). Researchers have used these signatures to develop statistical learning strategies for recognizing individuals within the same species (Stowell et al., 2019; Bedoya and Molles, 2021a; Huang et al., 2025a; Chen et al., 2020). However, most of these strategies are tailored to species with short and stereotyped vocalizations. This assumption presents a significant limitation when applied to species with prolonged or complex songs, as fixed-size input strategies often fail to leverage information distributed across extended vocalizations for accurate identification. Therefore, models capable of handling variable-length vocalizations and capturing long-range temporal patterns are needed.

Despite progress toward automated acoustic individual identification (AIID), most current bioacoustic systems are optimized for species-level recognition (Knight et al., 2024). Early attempts of AIID relied on handcrafted features and classical algorithms. For instance, Budka et al. (2015) used pulse-to-pulse duration with discriminant analysis to achieve 98% accuracy for 122 male corncrakes (*Crex crex*). Similarly, Ptacek et al. (2016) employed Gaussian mixture models with a universal background model (GMM-UBM) on linear frequency cepstral coefficients (LFCCs), reaching 78.5% accuracy for 13 chiffchaff males (*Phylloscopus collybita*). However, these promising results were confined to species with highly stereotyped vocalizations (low variability) and demanded extensive per-individual training data.

More recently, deep-learning and data-augmentation strategies have mitigated some of these limitations. Stowell et al. (2019) combined features learned by a convolutional neural network (CNN) with a random forest (RF) classifier, surpassing 85% area under the ROC curve (ROC-AUC) across 16 little owls (*Athene noctua*), 23 chiffchaffs, and 20 tree pipits (*Anthus trivialis*). Building on this work, Huang et al. (2025a) introduced AemNet, a 1-D CNN designed to process raw audio. They evaluated this architecture on a subset of the aforementioned data, as well as on 16 red-tailed black cockatoos (*Calyptorhynchus banksii graptogyne*) and 15 little penguins (*Eudyptula minor*), achieving accuracies of 92–97%. Additionally, Bedoya and Molles (2021a) integrated a CNN with the fuzzy clustering algorithm LAMDA to identify 10 male and 10 female great-spotted kiwis (*Apteryx maxima*) with 89–95% accuracy. Despite this progress in AIID, these models often overlook long-range temporal patterns and still require large per-individual training datasets, limiting their scalability to species with lengthy or variable songs.

Transfer learning, defined as repurposing broadly trained models to extract feature representations (embeddings) for new, data-sparse tasks (Weiss et al., 2016), has recently aided large-scale biodiversity monitoring (Stowell, 2022) and holds strong potential for individual-level identification (Knight et al., 2024). For example, embeddings from publicly available bird-classification models, such as BirdNET (Kahl et al., 2021), Perch (Google, 2023), and HawkEars (Huus et al., 2025), have been successfully repurposed to identify vocalizations when species identity is unknown (McGinn et al., 2023), detect novel sound classes (Allen-Ankins et al., 2025), and even classify non-avian taxa (Ghani et al., 2023). A pertinent application of transfer learning to individual identification is the work by Merino Recalde et al. (2025), which built on the densely sampled and richly annotated great tit song dataset described by Merino Recalde et al. (2024). They processed songs from over 400 great tit (*Parus major*) individuals using a pretrained Vision Transformer (ViT-S/16) on spectrogram inputs. The resulting embeddings were mapped into a similarity space that captures individual-specific acoustic cues. Their system achieved a Cumulative Matching Characteristic at rank 1 (CMC@1) of 98%, which indicates the percentage of queries where the correct individual was the highest-ranked prediction. Furthermore, they reported a mean Average Precision at 5 (mAP@5) of 98%, a metric that evaluates the precision and ordering of correct matches within the top five results, demonstrating highly robust retrieval performance.

Building on these applications, embeddings generated by bioacoustic foundation models provide trans-ferable representations that can be adapted to individual-identification pipelines, even with limited labeled data (Kath et al., 2024; Schwinger et al., 2025). These models typically process fixed-duration audio segments (e.g., 3 to 10 s), converting spectrogram inputs into fixed-dimensional embeddings via a convolutional backbone (Kahl et al., 2021). However, because this approach treats audio segments in isolation, it inherently discards the long-range temporal structure of complex bird songs. This limitation suggests that sequence-aware methods, capable of modeling the dependencies between successive embedding windows, could significantly improve individual-level identification.

One strategy for modeling these temporal patterns is to use recurrent neural networks (RNNs). Unlike models that treat each audio segment independently, RNNs process sequences by maintaining a hidden state that summarizes prior inputs (Goodfellow et al., 2016b). This hidden state can be viewed as a running summary of the recent context in the embedding sequence, enabling the model to capture dependencies across consecutive embedding windows that would be invisible if each window were analyzed in isolation. Consequently, the RNN can encode how note durations, inter-note intervals, and their syntactic arrangement evolve within a vocal sequence, which may carry individual identity. However, classical RNNs struggle with long-range dependencies due to exploding and vanishing gradients (Pascanu et al., 2013). Therefore, the standard approach to mitigate this issue is to employ long short-term memory (LSTM) or gated recurrent unit (GRU) networks, which utilize gating mechanisms to regulate information flow (Graves, 2012; Dey and Salem, 2017).

While recurrent architectures are widely used in sequential domains such as speech recognition and machine translation (Graves et al., 2013; Bahdanau et al., 2016), their use in bioacoustics has focused mostly on species-level classification (Gong et al., 2021; Makropoulos et al., 2023). A recent CNN–BiLSTM approach for bird species recognition further illustrates the value of recurrent temporal modeling for capturing long-range vocal dynamics (Sharma et al., 2026). However, this approach addresses species-level recognition and trains a task-specific CNN–recurrent architecture from spectrotemporal representations, rather than combining pretrained bioacoustic embeddings with recurrent modeling for individual-level identification. To our knowledge, no study has integrated large-scale acoustic embeddings from pretrained models with recurrent architectures to model intra-individual temporal structure. We hypothesize that processing time-ordered window-level embeddings from pretrained bioacoustic models via an LSTM offers a scalable approach for individual identification. This method is intended to leverage both short-window spectrotemporal information and sequence-level context, particularly for species with extended or variable songs.

In this study, we introduce a framework for individual-level bird identification based on vocalization sequences. Specifically, our approach uses an LSTM network to process sequences of time-ordered bioacoustic embeddings extracted via transfer learning from BirdNET. BirdNET V2.4 was selected due to its public availability, broad taxonomic coverage, and its consistently strong performance using shallow finetuning, particularly in complex acoustic environments (Ghani et al., 2024). We evaluate the approach on multiple publicly available datasets previously used for individual recognition, enabling comparison with established methods based on classical classifiers, CNNs, and vision transformers. By applying sequence modeling to learned embeddings, the architecture mitigates one limitation of fixed-window approaches by allowing multiple consecutive embedding windows to contribute to the identification decision, particularly for species with extended or variable song structures. Specifically, we pursue three aims: (i) evaluate whether processing sequences of transfer-learned embeddings improves individual-level identification compared with fixed-size static summaries; (ii) assess how vocalization duration, chronological order of embeddings, and pretrained representation choices affect identification performance; and (iii) evaluate the robustness of the pipeline across species, subset definitions, and recording conditions without relying on data augmentation.

The remainder of this article is organized as follows: we first describe the datasets and then detail the proposed two-stage pipeline, comprising embedding extraction and recurrent classifier construction. We then present the experimental setup, including the repeated-seed evaluation protocol, summarize the baseline and ablation experiments, and analyze the results relative to previous studies. Finally, we discuss the implications, limitations, and practical relevance of the proposed framework.

## 2. MATERIALS AND METHODS

### 2.1. Datasets

We evaluated our framework on seven datasets, each representing a distinct bird population and organized into within-year or across-year subsets when available. The nine subsets were defined following the original studies’ conventions, separating recordings collected within a single breeding season from those spanning multiple years. These datasets have been used in individual-recognition research and derive from four publicly available collections of labeled vocalizations: (i) the compilation by Stowell et al. (2018), which includes three species (chiffchaff, little owl, and tree pipit); (ii) the great spotted kiwi dataset of Bedoya and Molles (2021b); (iii) a collection of red-tailed black cockatoo and little penguin vocalizations (Huang et al., 2025b); and (iv) the Wytham great tit song dataset, available through OSF and described in detail as a richly annotated acoustic resource (Merino Recalde, 2023; Merino Recalde et al., 2024).

All recordings were obtained using autonomous recording units or handheld devices in natural habitats, following the protocols described in the original publications. Sampling rates ranged from 8.0 to 48.0 kHz, with 16-bit depth; all files were provided as single-channel audio. Only vocalizations with unambiguous individual identity, confirmed through field observations or tagging, were retained for this study. The only exception was the little penguin subset, in which vocalizations were labeled by nest rather than by individual, resulting in a nest-level classification task. This last subset was included to evaluate whether the framework can operate under weaker identity resolution. In total, the study comprises 87,865 audio segments of labeled vocalizations from 352 individuals.

These datasets cover a range of vocalization types, from short and stereotyped calls to long, variable songs, allowing us to assess model performance under diverse acoustic conditions. Datasets with *high complexity* typically involved a larger number of individuals and/or greater variability in vocalization syntax, while *low-complexity* datasets exhibited fewer individuals and more stereotyped vocal patterns. Together, these datasets offer a representative range of species whose vocal signatures have been previously shown to contain individual-specific acoustic cues.

Table 1 presents a summary of the datasets used in this study. For each species, we report the recording parameters (sampling rate and mean duration) and dataset statistics, including the number of labeled individuals, total vocalizations, and the variability of vocalizations per individual (min, max, and mean ± SD). Representative spectrograms of vocalizations per species are shown in Fig. 1, illustrating the differences in temporal and spectral structure.

**Table 1:**
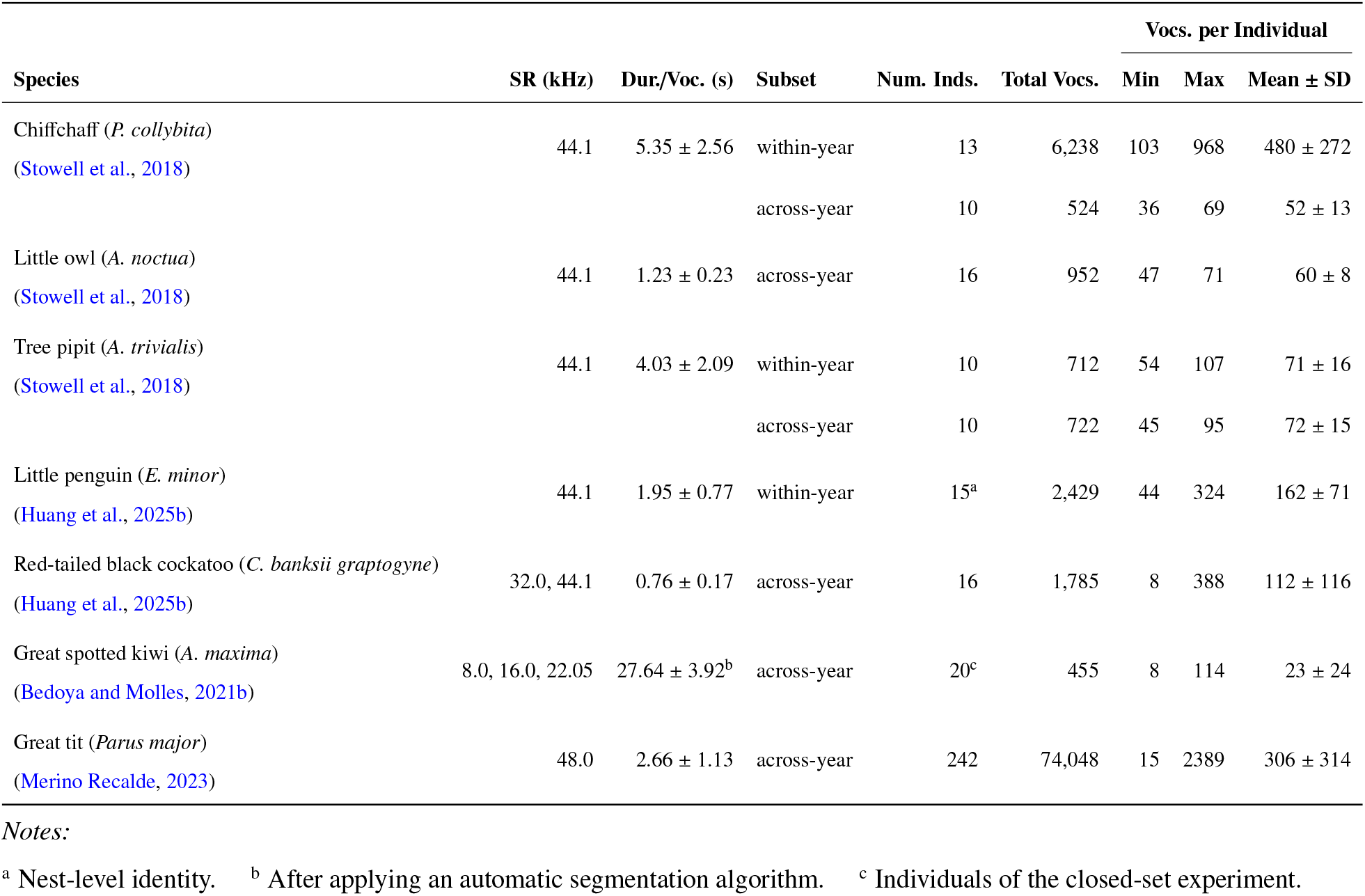
Summary of datasets employed in this study. Columns detail the recording sampling rate (SR), vocalization duration (Dur./Voc.), subset definition, total number of vocalizations (Total Vocs.), and class distribution statistics: number of individuals (Num. Inds.) and vocalizations per individual (Vocs. per Individual). Values for Dur./Voc. are reported as mean ± SD, while Vocs. per Individual includes min, max, and mean ± SD.

**Figure 1:**
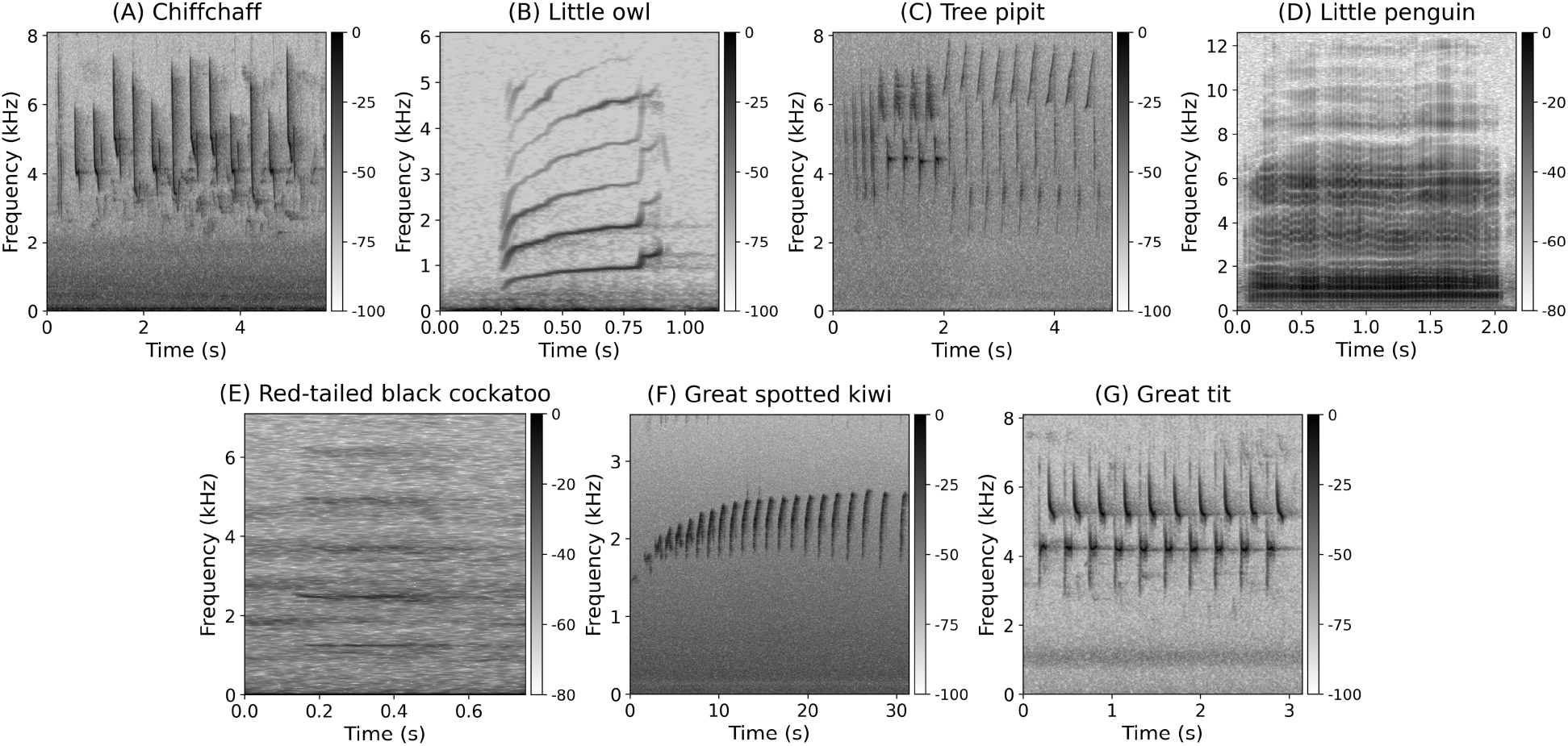
Representative spectrograms of the seven species included in this study. The figure illustrates the structural diversity of the vocalizations, ranging from short calls to complex songs. The panels show: (A) Chiffchaff, (B) Little owl, (C) Tree pipit, (D) Little penguin, (E) Red-tailed black cockatoo, (F) Great spotted kiwi, and (G) Great tit. Note the varying time scales (x-axis) and frequency ranges (y-axis) reflecting the natural characteristics of each species. All spectrograms were generated using a 2048-point STFT with a Hanning window and 75% overlap (hop length of 512 samples).

### 2.2. Proposed framework

In this work, we developed a framework that combines transfer learning with recurrent neural networks (specifically LSTMs) for individual identification based solely on acoustic signals. The system takes as input a set of vocalizations previously segmented in time (with defined onset and offset) and labeled by individual. Given that the primary objective is to evaluate the ability of temporally concatenated transfer-learned embeddings to improve individual identification on pre-processed data, this study does not consider earlier stages such as noise attenuation or acoustic event detection. Consequently, we utilized the segmented audio datasets described in Table 1, as they meet these criteria. However, for the great spotted kiwi dataset, we applied an additional segmentation algorithm since the original segments were 60 s long and contained background noise surrounding the actual vocalization; details are provided in Supporting Information S1. An overview of the complete workflow used to design the classification framework is presented in Fig. 2 and is explained in the sections below.

**Figure 2:**
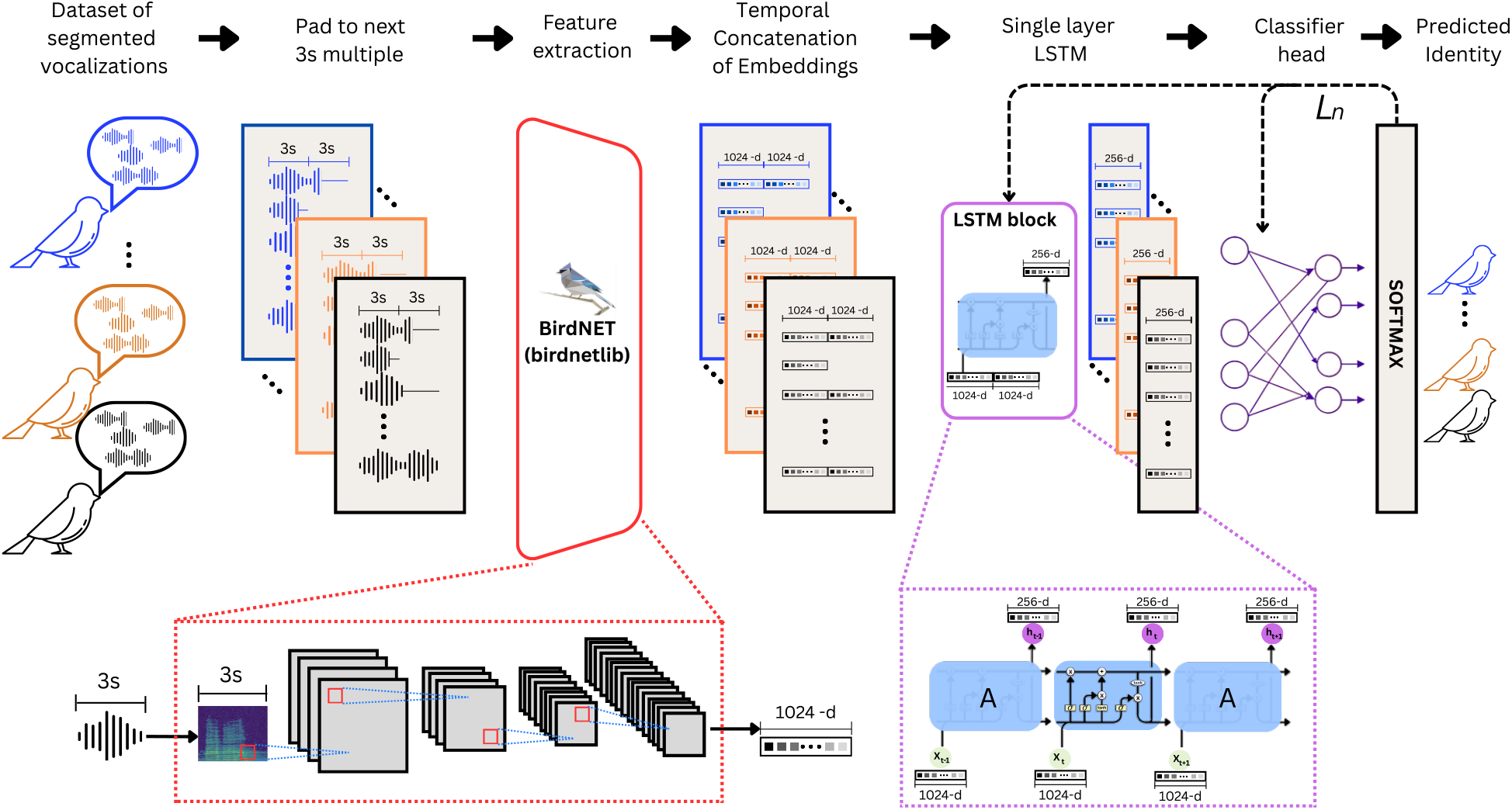
Architecture of the proposed acoustic identification system. Segmented vocalizations are padded to the next 3 s multiple and processed by BirdNET to extract 1024-dimensional embeddings. These embeddings are temporally concatenated and fed into a single-layer LSTM with 256 hidden units. The classifier head uses a Softmax activation to determine the predicted identity. The model is optimized using Cross-Entropy Loss (*L*_*n*_).

#### 2.2.1. Pre-processing and Embedding Extraction

The first stage of the methodology applied zero-padding to the end of each segmented vocalization until its duration reached the next multiple of 3 s. This was applied only when the duration of a segmented vocalization was not an exact multiple of 3 s. This temporal normalization ensures compatibility with BirdNET V2.4, which processes audio in fixed 3 s windows. We implemented this step using the `librosa` (McFee et al., 2015) and `soundfile` (Bechtold and Geier, 2023) libraries. We then divided each vocalization into non-overlapping 3 s windows, and all resulting windows were retained for embedding extraction. No window-level energy threshold was applied to discard low-information or padding-dominated windows after this padding step. We extracted features using the BirdNET V2.4 backbone without the final classification layer, yielding a 1024-dimensional embedding for each window. The feature extraction stage relies on `birdnetlib` (Weiss, 2023), which facilitates integrating BirdNET-Analyzer functionality into Python environments. It automates key preprocessing steps, including resampling audio to 48 kHz and generating the spectrograms required by the model.

Consequently, the number of generated embeddings varies depending on the total duration of the padded vocalization. To preserve the temporal structure, we arranged the embeddings from each window in chronological order. In this way, each vocalization is represented by a time-ordered sequence of *n* embeddings, where *n* equals the padded duration divided by the 3 s window length. For instance, a vocalization padded to 9 s results in a sequence of three embeddings (*n* = 3). No data augmentation techniques, such as time-stretching, pitch-shifting, or background noise injection, were applied. This constraint was strictly enforced to evaluate the intrinsic discriminative power of the pretrained embeddings and the proposed temporal modeling architecture without the confounding influence of synthetic data variability.

#### 2.2.2. Recurrent Classification Network

Subsequently, the concatenated sequences of variable-length embeddings were fed into a single-layer LSTM network (see Graves (2012) for a comprehensive explanation of LSTM). Because vocalizations generated different numbers of BirdNET embeddings, sequences within each mini-batch were padded at the tensor level and then processed as packed sequences. This allowed the LSTM to use only valid embedding time steps while ignoring artificial batch-padding positions. This procedure is distinct from the audio zero-padding applied before BirdNET extraction and does not remove low-information audio windows. The LSTM treats the input structure as a time series, processing the temporal information from each vocalization. Through its final hidden state, the network summarizes each sequence into a single embedding with a dimensionality of 256. These embeddings, which combine short-window spectrotemporal information with sequence-level context, are finally passed to a fully connected neural network with a softmax activation function to estimate class membership probabilities.

### 2.3. Experimental setup

To evaluate the proposed framework, each of the nine subsets was evaluated over five independent random seeds. For each seed, the data were partitioned into training (60%), validation (20%), and test (20%) sets using a stratified split strategy. This approach ensured that all individuals (classes) were represented across all partitions, while strictly maintaining that the specific vocalizations used for testing were different from those in the training and validation sets. The same seed-specific partitions were used for all model configurations evaluated within a given subset, enabling paired comparisons across architectures. This protocol allowed us to evaluate generalization to unseen vocalizations of known individuals and to quantify the sensitivity of the reported metrics to random data partitioning. However, these stratified vocalization-level splits do not necessarily correspond to session-disjoint evaluations. Consequently, the reported metrics should be interpreted as generalization to unseen vocalizations of known individuals under repeated stratified partitioning, rather than as strict re-identification across independent recording sessions.

Given the classification task, the model was trained by minimizing the standard cross-entropy loss function (Goodfellow et al., 2016a). The training process was optimized using the Adam algorithm (Kingma and Ba, 2015) with a constant learning rate of 0.001. To stabilize recurrent layer training, gradient clipping with a threshold of 1.0 was applied. The model was trained for a maximum of 40 epochs with a batch size of 32, and early stopping was implemented with a patience of 10 epochs. Crucially, to mitigate the potential impact of class imbalance during model selection, we monitored the macro-averaged F1-score on the validation set to select the best checkpoint, ensuring that performance reflected the model’s ability to identify both frequent and rare individuals while also preventing overfitting.

To characterize failure cases of our proposed framework, we conducted an illustrative per-individual error analysis on the great spotted kiwi across-year subset using the ordered BirdNET + LSTM model across the five random seeds. This subset was selected because it contains long vocalizations and showed one of the clearest gains from recurrent aggregation. For each seed, we used the test-set predictions and seed-specific split assignments to compute the number of errors per individual, per-individual recall, training-set size per individual, recurrent confusion pairs, and Spearman correlations between training-set size and both recall and error rate (see Supporting Information S3).

For each subset, seed, and model configuration, training was performed independently using the same optimization protocol. Unless otherwise stated, our reported results correspond to the mean ± standard deviation across the five seeds. As a limited sensitivity check on the great spotted kiwi across-year subset, we evaluated LSTM hidden sizes of 64, 128, 256, and 512 units with a single fixed seed, keeping all other hyperparameters unchanged. The 256-unit configuration achieved the highest macro-F1, with the 512-unit configuration yielding similar but slightly lower performance; therefore, we retained 256 hidden units for all experiments.

All experiments were conducted using the PyTorch Lightning framework (Falcon and The PyTorch Lightning Team, 2019) in a standard cloud-based environment (Lightning AI Studio) with 4 vCPUs, without GPU acceleration. To provide a quantitative reference for CPU-only deployment, we measured post-segmentation inference time. Because the proposed workflow operates on pre-segmented vocalizations rather than continuous audio streams, runtime was benchmarked on a representative great spotted kiwi vocalization with an original duration of 27.624 s, which was padded to 30 s and therefore corresponded to 10 non-overlapping BirdNET windows. Since BirdNET embedding extraction and LSTM classification were implemented in two decoupled computational environments, these stages were benchmarked separately. The BirdNET stage was repeated 20 times with the analyzer already loaded, and runtime was recorded for audio loading, zero-padding, temporary file writing, and embedding extraction. LSTM classification was measured over 1000 repetitions using the exported BirdNET embeddings and the trained model loaded in CPU evaluation mode. The total post-segmentation inference time was estimated by summing the mean runtimes of the measured stages.

### 2.4. Baseline and ablation experiments

To rigorously evaluate the modeling assumptions underlying the proposed framework, we designed a set of baseline and ablation experiments. These experiments addressed three related questions: (i) whether individual identity can be inferred from fixed-size summaries of pretrained embeddings, rather than from a sequence of embeddings; (ii) whether preserving the chronological order of embeddings improves performance beyond the nonlinear aggregation capacity of the LSTM; and (iii) how an alternative pretrained representation coupled with static downstream classifiers performs relative to the proposed recurrent BirdNET-based configuration. An overview of all baseline and ablation pipelines and controlled variables is shown in Fig. 3.

**Figure 3:**
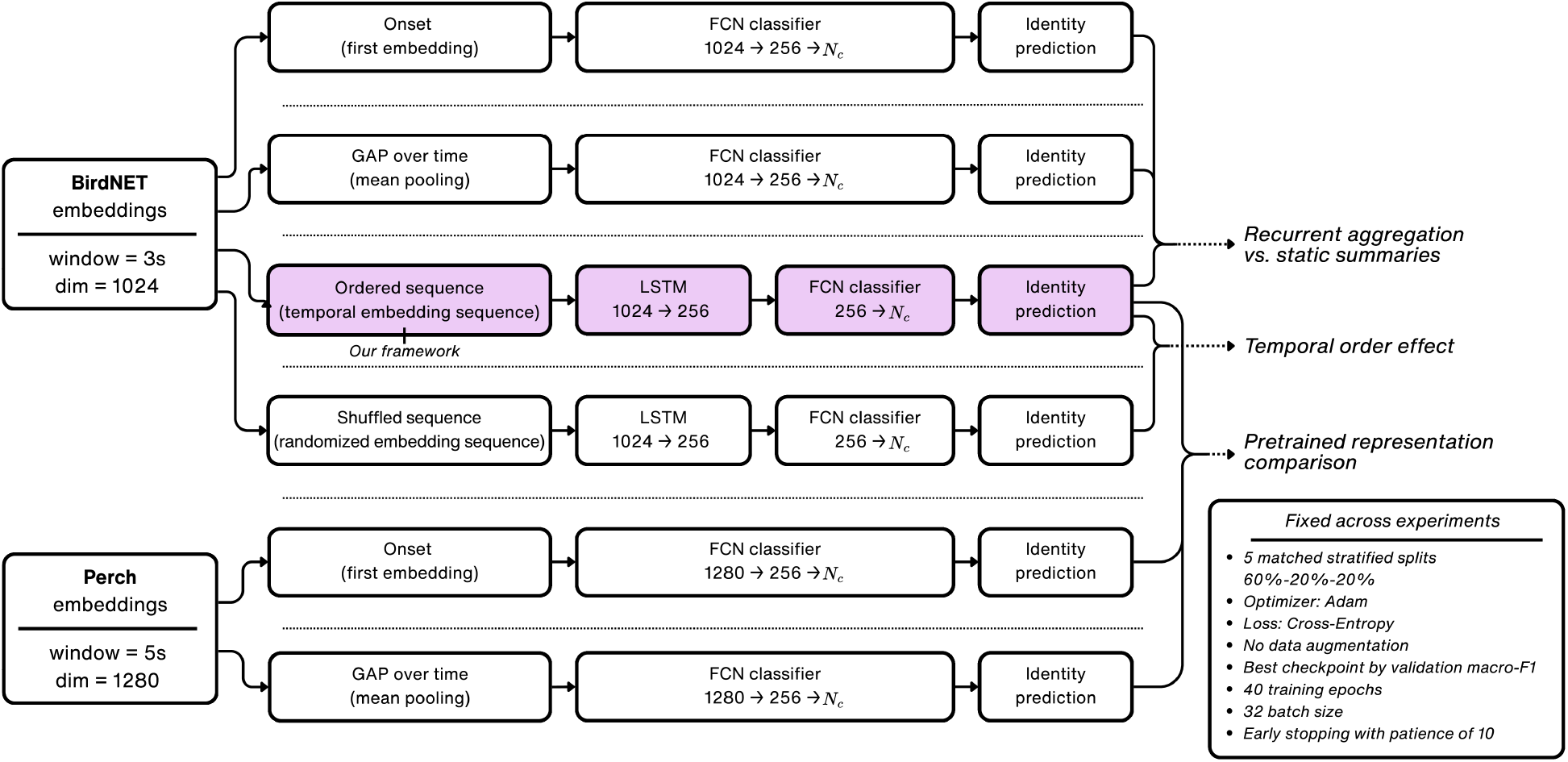
Schematic of the baseline and ablation experiments. The diagram summarizes three comparisons: recurrent aggregation versus static summaries, ordered versus shuffled LSTM sequences, and comparisons of pretrained representations. BirdNET embeddings (3 s windows, 1024D) are evaluated using (i) an Onset-only fully connected network (FCN) baseline, (ii) a global average pooling (GAP) FCN baseline, (iii) the proposed ordered LSTM framework (highlighted in light purple), which consists of a chronological embedding sequence processed by an LSTM (1 layer, hidden=256) followed by an FCN classifier, and (iv) a shuffled-order LSTM ablation, in which the same embeddings are randomly permuted within each vocalization before being processed by the same LSTM architecture. Perch embeddings (5 s windows, 1280D) are evaluated with non-recurrent FCN baselines (Onset-only and GAP) to compare an alternative pretrained bioacoustic representation under static downstream classifiers. In all architectures, *N*_*c*_ denotes the number of individuals (classes). Brackets highlight the intended comparisons without assuming that any single comparison isolates all sources of variation. The bottom box summarizes experimental settings held fixed across all runs to ensure a fair comparison.

Crucially, these controlled ablations serve as a sensitivity analysis of key modeling assumptions in our workflow. By varying only one component at a time (temporal order, recurrent aggregation versus static summaries, and pretrained representation choice) while holding data splits and training protocol fixed, we quantify how sensitive AIID performance is to these design choices across species and recording conditions.

#### 2.4.1. Recurrent sequence modeling versus static baselines

We evaluated the contribution of the recurrent architecture by replacing the LSTM layer with a fully connected network (FCN). This comparison helps determine whether individual identification benefits from processing a sequence of embeddings, rather than representing each vocalization through a single fixed-size vector. However, unlike LSTMs, FCNs require fixed-size inputs. Consequently, we adopted two complementary strategies to handle variable-length embedding sequences:

- *Onset-based baseline*: The input to the classifier was restricted to the first embedding vector of each sequence, corresponding to the initial 3 s of the vocalization. This baseline evaluates whether individual identity is sufficiently encoded in the vocalization onset alone, which is particularly relevant for species characterized by short, abrupt calls.
- *Pooling-based baseline*: Global average pooling (GAP) aggregates all embeddings within a sequence into a single mean vector. By applying GAP across the temporal dimension, we explicitly discard temporal-order information while retaining information from all embedding windows. This approach tests whether identity can be resolved using only the global short-term spectrotemporal summary of the vocalization, without explicitly modeling the order of the embeddings. Such a baseline is particularly relevant for species with sustained or highly repetitive vocal patterns. In these cases, a global summary of the available embeddings may be sufficient for individual discrimination.

For species with short vocalizations (< 3s), which typically result in a single embedding per vocalization (*n* ≈ 1), the Onset and GAP baselines are expected to converge because both operate on the same embedding. These cases provide a reference condition for the LSTM when little or no temporal sequence is available to model, and therefore are not used to infer the contribution of temporal ordering. In contrast, for species with longer, acoustically complex vocalizations (*n* > 1), the comparison between the GAP baseline and the LSTM-based model assesses whether recurrent aggregation improves performance beyond simple averaging of the available embeddings. This ablation design explicitly accounts for the heterogeneity in vocalization duration across the seven target species.

To quantify the contribution of the LSTM framework relative to the static baselines, we define the mean signed difference 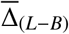 for each species and subset as:

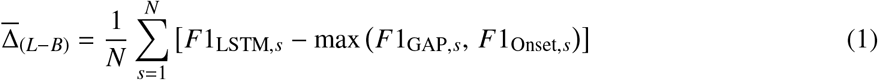

where *N* = 5 is the number of random seeds and max (*F*1_GAP,*s*_, *F*1_Onset,*s*_) denotes the stronger of the two static baselines at seed *s*. A positive value indicates that the LSTM outperforms the strongest static baseline on average. To assess whether 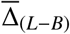 differs significantly from zero, we applied a paired *t*-test across the five seeds, using significance thresholds of α = 0.05 and α = 0.01.

#### 2.4.2. Ordered versus shuffled LSTM ablation

The comparison between the LSTM and the GAP baseline evaluates whether recurrent aggregation improves performance beyond simple averaging of the embedding sequence. However, this comparison does not fully isolate the effect of temporal order, because the LSTM may also benefit from nonlinear and adaptive aggregation of multiple embeddings. To disentangle the contribution of chronological order from the LSTM’s nonlinear aggregation capacity, we implemented an additional shuffled-order LSTM ablation.

In this experiment, the same BirdNET embeddings were provided to the LSTM, but the order of embeddings within each vocalization was randomly permuted. Specifically, after embeddings were sorted chronologically by window start time, a seed-specific random permutation was generated once for each vocalization and then kept fixed throughout training, validation, and testing. This condition preserves the number of embeddings, the embedding content, the recurrent architecture, and the train/validation/test partitions used by the ordered LSTM, while disrupting the original chronological arrangement of the sequence. Vocalizations represented by a single embedding remained unchanged because no temporal order can be permuted. Thus, a performance reduction in the shuffled condition would support the interpretation that chronological order contributes information beyond the mere availability of multiple embedding windows.

We quantified the effect of preserving chronological order using the paired mean signed difference 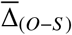, defined analogously to Eq. (1) as the test macro-F1 of the ordered LSTM minus the test macro-F1 of the shuffled LSTM for the same seed. A positive value indicates that preserving the chronological order of embeddings improves performance. We assessed whether 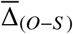 differed from zero using a paired *t*-test across the five random seeds.

#### 2.4.3. Comparison of pretrained feature representations

We evaluated an alternative feature representation using Perch (Google, 2023) to compare two widely used pretrained bioacoustic embedding models under the same downstream classification framework. Unlike BirdNET V2.4, which extracts 1024-dimensional embeddings from 3 s segments, Perch processes 5 s audio windows and produces 1280-dimensional embeddings. Perch embeddings were coupled exclusively with the fully connected classifier strategies described above (Onset-based and Pooling-based), rather than with an LSTM, so that the comparison remained focused on non-recurrent transfer-learning baselines. Under this design, observed performance differences reflect the combined effects of the pretrained representation, embedding dimensionality, training background, and input-window configuration.

### 2.5. Training protocol and fair comparison

To ensure a fair and controlled comparison across all models, we adhered to the same seed-specific train/validation/test partitions, initialization scheme, and training protocol detailed in subsection 2.3 for all experiments. Specific preprocessing adjustments were applied when using Perch, such as padding audio segments to the next multiple of 5 s to match its native window size. Regarding model architecture, the input layer of the fully connected baselines was instantiated to match the native embedding dimension of the respective backbone (1024 for BirdNET and 1280 for Perch). Crucially, the hidden layer size was fixed at 256 units across all experiments, matching the hidden dimensionality of the LSTM used in the main framework. This design choice reduces differences in downstream classifier capacity and supports controlled comparisons within the constraints of each pretrained representation.

### 2.5. Comparison with state-of-the-art approaches

To benchmark the proposed framework against existing AIID systems, we compare our results with previously published metrics reported on the same datasets. Since we do not reimplement external models, comparisons are limited to studies whose dataset definitions and task settings are compatible with ours (e.g., within-year vs. across-year subsets, and closed-set identification). However, we explicitly distinguish between subset-level compatibility and protocol-level equivalence. Some previous benchmarks, particularly those based on the datasets compiled by Stowell et al. (2018), used day-held-out or year-held-out splits, whereas our evaluation uses repeated stratified vocalization-level splits. Therefore, when split protocols differ, we report the closest compatible subset, explicitly note the mismatch, and interpret these comparisons as contextual benchmarks rather than direct evidence of superiority under the original evaluation design.

For chiffchaff, little owl, and tree pipit (from the collection of Stowell et al. (2018)), we compare against two prior approaches: the CNN-feature coupled with a random forest pipeline of Stowell et al. (2019) and the raw-audio AemNet model of Huang et al. (2025a). For red-tailed black cockatoo and little penguin, comparisons are based on the results reported by Huang et al. (2025a). For great spotted kiwi, we benchmark against the closed-set identification experiment of Bedoya and Molles (2021a), which matches our task definition of recognizing known individuals. For the great tit, we compare against the ViT-based deep metric learning framework used by Merino Recalde et al. (2025) for song-based re-identification across time, which evaluates identification as a ranking problem. This provides a relevant benchmark because the study applies acoustic re-identification to a large wild population.

Considering that reported metrics vary across studies, each comparison is aligned to the metrics explic-itly provided in the corresponding reference. For Stowell et al. (2019), performance is reported primarily as ROC-AUC under multiple experimental assumptions; therefore, we cite the best-performing configuration per species as reported in that work. For AemNet (Huang et al., 2025a), we report the published ROC-AUC and accuracy values when their subset definition matches ours (notably, chiffchaff and tree pipit were evaluated only in the within-year setting). For great spotted kiwi, Bedoya and Molles (2021a) report sex-stratified accuracies for the closed-set experiment, separately for 10 males and 10 females. In contrast, our experiment used the same balanced closed-set composition at the individual level, but reports pooled accuracy without sex stratification. Therefore, our result should be interpreted as an overall closed-set estimate for the balanced 20-individual pool, rather than as directly comparable to either the male- or female-specific accuracy reported in the reference study.

For the great tit, Merino Recalde et al. (2025) report retrieval metrics (CMC@1 and mAP@5). To enable a direct comparison under the same ranking-based perspective, we compute CMC@1 and mAP@5 from our model’s class scores by treating each test vocalization as a query and ranking candidate identities by the predicted logits (top-*k*). Under this formulation, CMC@1 corresponds to top-1 accuracy in the ranked list, while mAP@5 summarizes whether the correct identity appears within the top five positions, with higher weight for higher ranks.

A key methodological distinction is that several published baselines considered here rely on data augmentation or task-specific training strategies, whereas our experiments intentionally avoid augmentation to isolate the discriminative power of pretrained embeddings and the evaluated downstream aggregation strategies, including recurrent sequence modeling. Thus, our results do not benefit from synthetic data expansion. However, because we rely on reported results, and because split protocols and preprocessing steps can differ across studies, these comparisons should be interpreted with caution. Some differences in preprocessing (e.g., segmentation conventions, filtering) may remain unavoidable; we therefore restrict comparisons to matching subsets whenever possible and indicate cases where only partial alignment is feasible.

## 3. RESULTS

### 3.1. Overall identification performance

The proposed framework, integrating BirdNET embeddings with a single-layer LSTM, achieved high identification performance across the seven bird species and nine experimental subsets. As summarized in Table 2, mean test macro-F1 scores ranged from 93.2±0.9% to 98.3±1.3%, while mean test accuracy ranged from 93.9 ± 1.7% to 98.3 ± 0.4% across the five random seeds. These results indicate that BirdNET embeddings coupled with LSTM-based sequence modeling provided strong discriminative performance across datasets with different vocalization durations, class distributions, and numbers of individuals. The framework maintained high identification accuracy for both frequent and underrepresented individuals without requiring loss weighting strategies.

**Table 2:** Overall performance of the proposed framework (BirdNET + LSTM) across the seven target species. Values are reported as mean ± standard deviation over five random seeds, with each seed defining the stratified train/validation/test partition and model initialization. Reported metrics include macro-averaged F1-score, accuracy, and area under the ROC curve (ROC-AUC). For species evaluated across multiple seasons, results are reported separately for within-year and across-year subsets. *N*_*c*_ denotes the number of individuals (classes) included in each subset-specific classification task; within-year and across-year rows are treated as separate evaluation scenarios when both are available.

| Species | Dur. (s)/Voc | Subset | $N_c$ | Macro-F1 (%) | Accuracy (%) | ROC-AUC |
| --- | --- | --- | --- | --- | --- | --- |
| Chiffchaff (*P. collybita*) | $5.35 \pm 2.56$ | within-year | 13 | $96.7 \pm 1.0$ | $97.5 \pm 0.8$ | $0.9996 \pm 0.0002$ |
| | | across-year | 10 | $98.2 \pm 0.4$ | $98.3 \pm 0.4$ | $0.9998 \pm 0.0001$ |
| Little owl (*A. noctua*) | $1.23 \pm 0.23$ | across-year | 16 | $98.3 \pm 1.3$ | $98.3 \pm 1.2$ | $0.9998 \pm 0.0003$ |
| Tree pipit (*A. trivialis*) | $4.03 \pm 2.09$ | within-year | 10 | $96.9 \pm 2.6$ | $97.1 \pm 2.4$ | $0.9995 \pm 0.0008$ |
| | | across-year | 10 | $95.2 \pm 3.4$ | $95.5 \pm 3.1$ | $0.9969 \pm 0.0029$ |
| Little penguin (*E. minor*) | $1.95 \pm 0.77$ | within-year | 15 | $93.2 \pm 0.9$ | $94.6 \pm 0.8$ | $0.9986 \pm 0.0006$ |
| Red-tailed black<br>cockatoo (*C. banksii graptogyne*) | $0.76 \pm 0.17$ | across-year | 16 | $95.9 \pm 1.2$ | $96.6 \pm 0.5$ | $0.9992 \pm 0.0006$ |
| Great spotted kiwi (*A. maxima*) | $27.64 \pm 3.92$ | across-year | 20 | $95.1 \pm 1.2$ | $93.9 \pm 1.7$ | $0.9982 \pm 0.0028$ |
| Great tit (*Parus major*) | $2.66 \pm 1.13$ | across-year | 242 | $94.0 \pm 0.5$ | $96.9 \pm 0.2$ | $0.9998 \pm 0.0001$ |

The model achieved high macro-F1 scores for the little owl and the chiffchaff across-year subset, reaching 98.3 ± 1.3% and 98.2 ± 0.4%, respectively. Notably, the framework maintained high performance in the most challenging scenario in terms of class number, the great tit dataset, achieving a macro-F1 score of 94.0 ± 0.5% despite the large number of individual classes (*N*_*c*_ = 242). For species represented by both within-year and across-year subset definitions, performance remained high under the repeated stratified split protocol. For example, the chiffchaff subsets reached macro-F1 scores of 96.7 ± 1.0% and 98.2 ± 0.4% for the within-year and across-year subsets, respectively. The lowest mean macro-F1 was observed in the little penguin nest-level classification (93.2 ± 0.9%), although performance remained high despite the weaker identity resolution of this subset.

In the CPU-only runtime benchmark, processing a representative great spotted kiwi vocalization padded to 30 s required 0.010 ± 0.001 s for audio loading, padding, and temporary file writing; 0.808 ± 0.016 s for BirdNET embedding extraction; and 0.0009 ± 0.0005 s for LSTM classification. This yielded a total post-segmentation inference time of 0.819 ± 0.016 s per 30 s padded vocalization. Most of the runtime was therefore associated with BirdNET embedding extraction, whereas the downstream LSTM classification step contributed only a very small fraction of the total inference time.

### 3.2. Effect of sequence modeling relative to static baselines

As shown in Table 3, the relationship between vocalization duration and the benefit of temporal sequence modeling was not strictly monotonic, although a general positive trend was observed. For this analysis, we report macro-F1 as the primary metric to reduce the influence of class imbalance. Additional performance metrics for all baseline configurations are provided in Supporting Information S2. The magnitude of the LSTM’s advantage over the stronger static baseline is quantified by 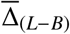 (see Eq. 1), and its significance was assessed via paired *t*-test across five random seeds.

**Table 3:**
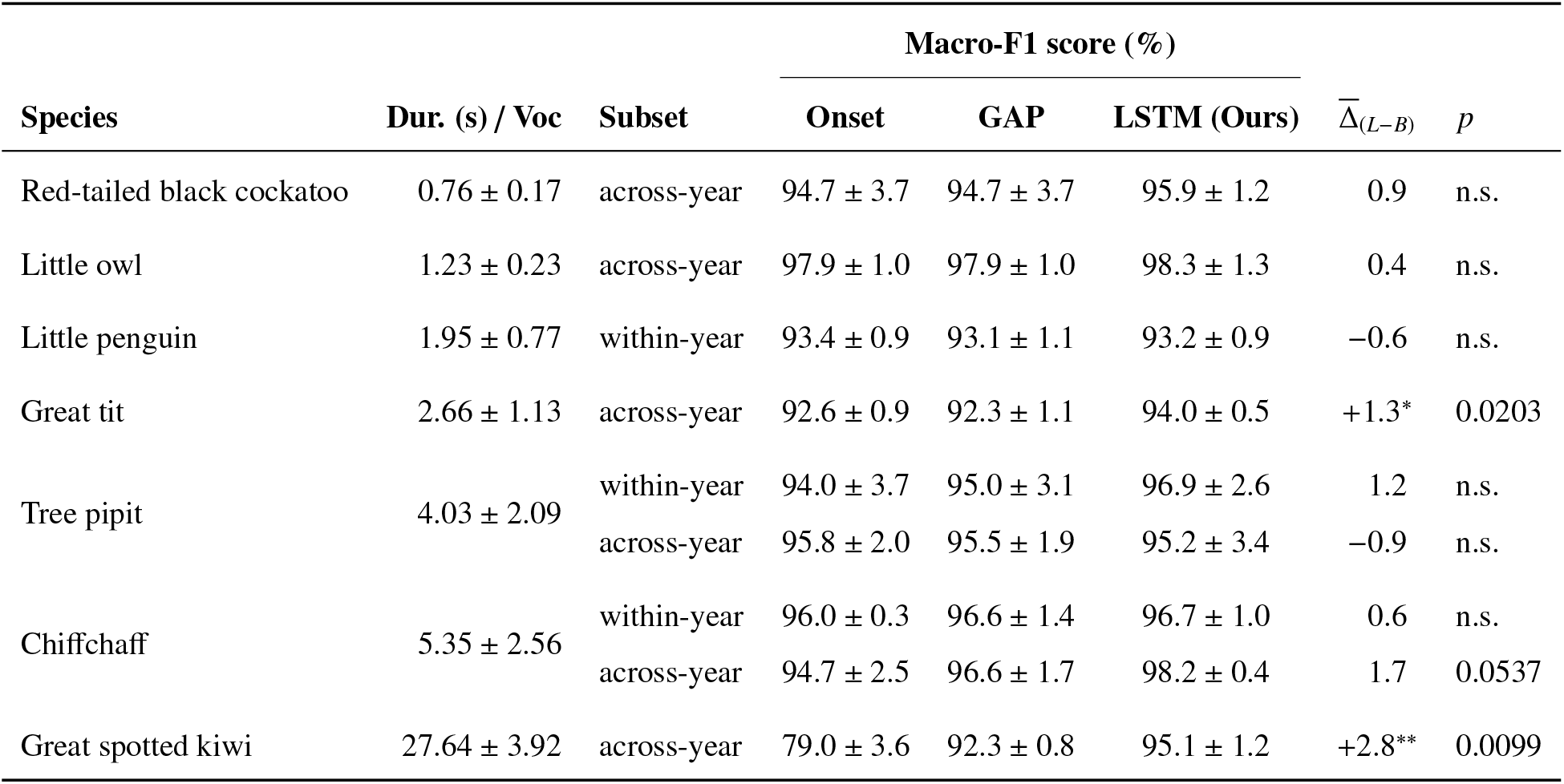
Ablation results for recurrent sequence modeling (BirdNET backbone) relative to static baselines ordered by mean vocalization duration. Performance is reported as mean ± standard deviation over five random seeds of the macro-F1 score (%). The 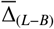 column reports the mean signed difference between the LSTM framework and the stronger static baseline at each seed. Statistical significance was assessed via a paired *t*-test. Markers: ^*^ *p*<0.05, ^**^ *p*<0.01.

Species with short vocalizations (<3 s), which typically yield a single embedding when processed with the BirdNET backbone, showed nearly identical macro-F1 scores across the three configurations. These high absolute scores should therefore be interpreted as evidence that BirdNET embeddings provided strong discriminative information for these datasets, rather than as evidence that the recurrent component alone drove performance. This pattern was observed for the red-tailed black cockatoo (0.76 ± 0.17 s) and the little owl (1.23 ± 0.23 s), for which the LSTM provided small, non-significant gains of +0.9 and +0.4 percentage points over the strongest static baseline, respectively. As expected, for both species, the Onset and

GAP baselines produced identical scores because pooling over a single embedding returns the same embedding. The little penguin (1.95 ± 0.77 s) followed a similar convergence pattern; here, the LSTM performed marginally below the best baseline 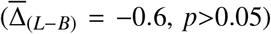, indicating that the recurrent architecture did not introduce a statistically detectable performance penalty when the temporal arrangement of embeddings was limited. Together, these three short-call species provide a reference case in which little or no temporal sequence is available to model. Their convergence is therefore expected and indicates no clear benefit or observed systematic degradation of recurrent modeling under single- or near-single-window conditions.

An exception to this short-call convergence was observed for the great tit (2.66 ± 1.13 s), where the LSTM achieved a statistically significant gain of +1.3 percentage points over the strongest static baseline (*p* = 0.0203), together with a reduction in inter-seed variance (SD = ±0.5 vs. ±0.9–1.1 for the baselines). Although the mean duration is close to the native 3 s analysis window of BirdNET, some great tit vocalizations exceeded this window and were represented by multiple embeddings. On average across the five seeds, approximately one-third of the test vocalizations generated more than one embedding. This provided the LSTM with limited but non-negligible sequence-level information, which may explain its advantage over the static baselines.

For the tree pipit (4.03 ± 2.09 s), results differed between evaluation subsets. The LSTM showed a positive but non-significant mean gain in the within-year condition 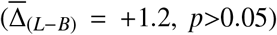, whereas it performed marginally below the best baseline in the across-year condition 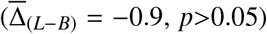. Thus, the high absolute performance observed for the tree pipit should also be interpreted as the performance of the BirdNET-based classification pipeline, rather than as clear evidence that the LSTM component provided a consistent advantage over static summaries. For the chiffchaff (5.35±2.56 s), the LSTM outperformed both baselines on average in both subsets, with a larger mean gain in the across-year condition 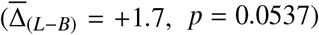 and a smaller non-significant gain in the within-year condition 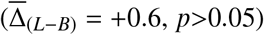.

The most pronounced benefits of sequential modeling were observed for the great spotted kiwi (27.64 ± 3.92 s). The Onset baseline lagged substantially behind GAP (79.0% vs. 92.3%, a difference of 13.3 percentage points), indicating that identity-relevant information was distributed across the full vocalization rather than concentrated at its onset. The LSTM framework further improved over GAP by 2.8 percentage points, with a statistically significant gain (*p* = 0.0099). Its inter-seed variance was lower than that of the Onset baseline (SD = ±1.2 vs. ±3.6), although slightly higher than that of GAP (SD = ±1.2 vs. ±0.8).

The illustrative error analysis for the great spotted kiwi subset (see Supporting Information S3) showed that misclassifications were not explained solely by the number of training vocalizations per individual. Across 455 test predictions from the five ordered-LSTM seeds, the model produced 28 errors. The association between mean training-set size and pooled recall was weak and non-significant (Spearman ρ = −0.211, *p* = 0.373), as was the association between mean training-set size and error rate (ρ = 0.211, *p* = 0.373). Instead, errors were partially concentrated in a small number of individuals: 11 of 20 individuals had at least one error, the three individuals with the most errors accounted for 60.7% of all errors, and the five individuals with the most errors accounted for 75.0%.

Across all species, the LSTM framework produced no statistically significant degradation relative to the strongest static baseline. Statistically significant improvements were observed only for the great tit and the great spotted kiwi. The latter represents the clearest case in which recurrent sequence modeling improved performance relative to static baselines, as it combined the longest vocalizations with the largest separation between the Onset and GAP baselines.

### 3.3. Comparison of pretrained feature representations (BirdNET vs. Perch)

We evaluated an alternative pretrained bioacoustic representation against the proposed BirdNET + LSTM framework. These results reflect the combined effect of the pretrained representation, its input-window configuration, and the downstream aggregation strategy. As shown in Table 4, the proposed Bird-NET + LSTM framework achieved higher macro-F1 than the Perch-based non-recurrent baselines across all species and subsets.

**Table 4:**
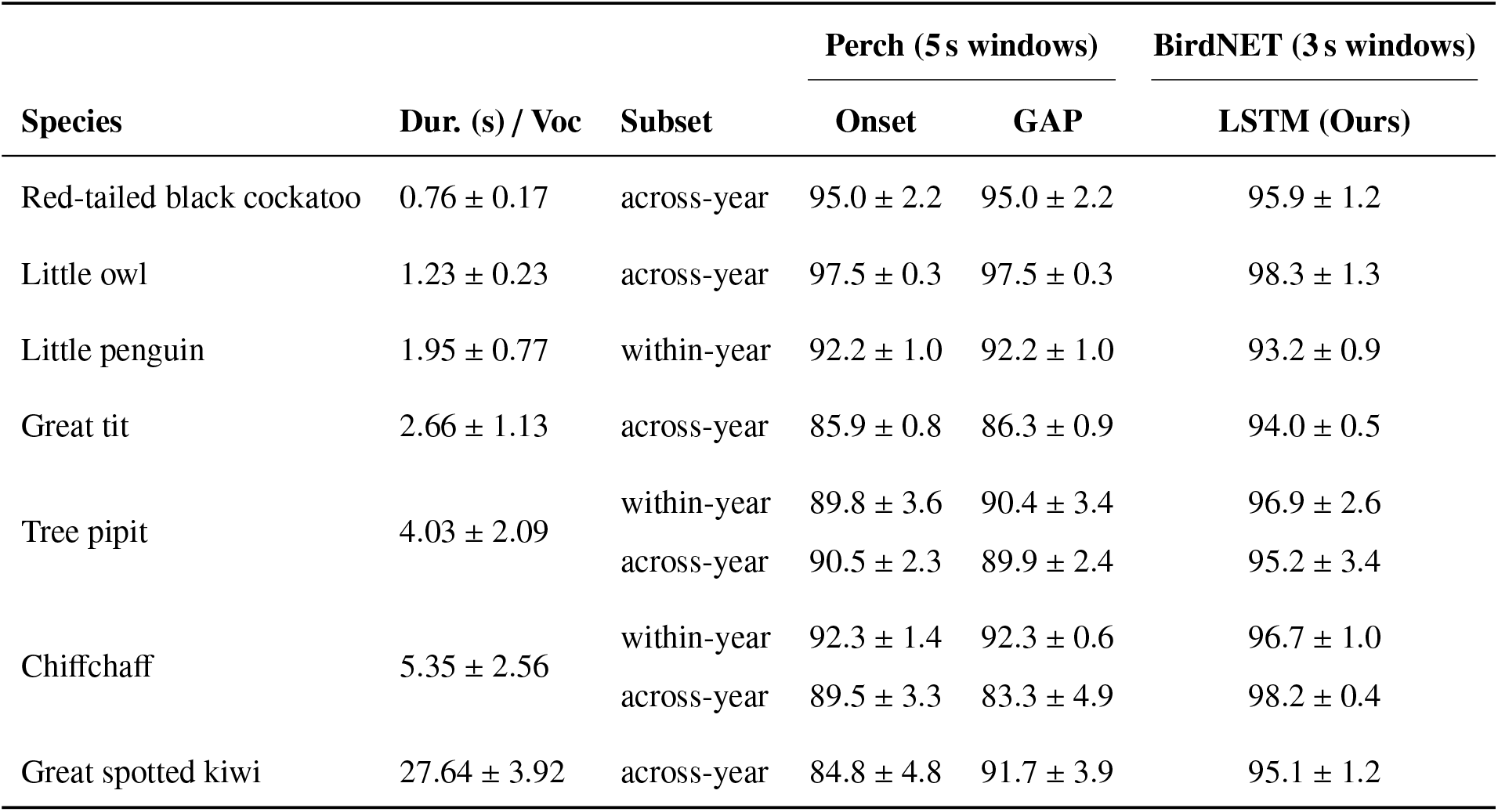
Comparison between Perch-based static baselines and the proposed BirdNET + LSTM framework. Performance is reported as mean ± standard deviation over five random seeds of the macro-F1 score (%). Perch embeddings (5 s windows, 1280D) are evaluated using Onset and GAP fully connected baselines, whereas the BirdNET column reports the proposed ordered LSTM framework using 3 s embeddings (1024D). This comparison contextualizes performance across pretrained bioacoustic representations and downstream aggregation strategies.

| Species | Dur. (s) / Voc | Subset | Perch (5 s windows) |  | BirdNET (3 s windows) |
| --- | --- | --- | --- | --- | --- |
|  |  |  | Onset | GAP | LSTM (Ours) |
| Red-tailed black cockatoo | 0.76 ± 0.17 | across-year | 95.0 ± 2.2 | 95.0 ± 2.2 | 95.9 ± 1.2 |
| Little owl | 1.23 ± 0.23 | across-year | 97.5 ± 0.3 | 97.5 ± 0.3 | 98.3 ± 1.3 |
| Little penguin | 1.95 ± 0.77 | within-year | 92.2 ± 1.0 | 92.2 ± 1.0 | 93.2 ± 0.9 |
| Great tit | 2.66 ± 1.13 | across-year | 85.9 ± 0.8 | 86.3 ± 0.9 | 94.0 ± 0.5 |
| Tree pipit | 4.03 ± 2.09 | within-year | 89.8 ± 3.6 | 90.4 ± 3.4 | 96.9 ± 2.6 |
| | | across-year | 90.5 ± 2.3 | 89.9 ± 2.4 | 95.2 ± 3.4 |
| Chiffchaff | 5.35 ± 2.56 | within-year | 92.3 ± 1.4 | 92.3 ± 0.6 | 96.7 ± 1.0 |
| | | across-year | 89.5 ± 3.3 | 83.3 ± 4.9 | 98.2 ± 0.4 |
| Great spotted kiwi | 27.64 ± 3.92 | across-year | 84.8 ± 4.8 | 91.7 ± 3.9 | 95.1 ± 1.2 |

The differences were small for the shortest vocalizations, where Perch Onset and GAP produced similar values and the BirdNET + LSTM advantage was below or close to one percentage point for the red-tailed black cockatoo, little owl, and little penguin. Larger numerical differences were observed for the great tit, tree pipit, chiffchaff, and great spotted kiwi. For example, in the great spotted kiwi, BirdNET + LSTM reached 95.1 ± 1.2%, whereas the best Perch baseline reached 91.7 ± 3.9%. These results indicate that, under the evaluated downstream configurations, the BirdNET-based framework provided stronger transfer-learning performance for AIID than the Perch-based static baselines. However, because the comparison combines differences in backbone representation and downstream model structure, it should not be interpreted as evidence that window duration alone explains the performance gap.

### 3.4. Effect of temporal order in LSTM embedding sequences

The ordered-versus-shuffled ablation was applied to the species and subsets for which temporal order could be meaningfully altered, that is, those containing vocalizations represented by more than one BirdNET embedding. As shown in Table 5, preserving the chronological order of embeddings did not produce a statistically significant improvement in most subsets. The only significant difference was observed for the chiffchaff across-year subset, where the ordered LSTM outperformed the shuffled LSTM by +1.9 percentage points (*p* = 0.0148).

**Table 5:** Effect of temporal ordering on BirdNET + LSTM performance for species with sufficient multi-embedding vocalizations to make shuffling informative. Results compare the original ordered embedding sequences against shuffled sequences. Performance is reported as mean ± standard deviation over five random seeds of the macro-F1 score (%). The 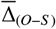 column reports the mean signed paired difference between the ordered and shuffled LSTM across matched seeds. Boldface indicates the best value between ordered and shuffled sequences. Statistical significance was assessed via paired *t*-test. Markers: ^*^ *p*<0.05.

| Species | Dur. (s) / Voc | Subset | Macro-F1 score (%) | | $\bar{\Delta}_{(O-S)}$ | $p$ |
| --- | --- | --- | --- | --- | --- | --- |
|  |  |  | Ordered LSTM | Shuffled LSTM |  |  |
| Great tit | 2.66 ± 1.13 | across-year | 94.0 ± 0.5 | 93.7 ± 0.5 | 0.3 | n.s. |
| Tree pipit | 4.03 ± 2.09 | within-year | 96.9 ± 2.6 | 95.4 ± 2.6 | 1.6 | n.s. |
| | | across-year | 95.2 ± 3.4 | 95.1 ± 1.3 | 0.1 | n.s. |
| Chiffchaff | 5.35 ± 2.56 | within-year | 96.7 ± 1.0 | 96.8 ± 0.4 | 0.0 | n.s. |
| | | across-year | 98.2 ± 0.4 | 96.3 ± 1.2 | +1.9* | 0.0148 |
| Great spotted kiwi | 27.64 ± 3.92 | across-year | 95.1 ± 1.2 | 92.5 ± 3.8 | 2.6 | n.s. |

Although statistical significance was limited to one subset, the largest mean differences were observed in species or subsets with longer vocalizations. The effect was small for the great tit 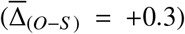, whose mean duration is close to the native 3 s BirdNET window, and negligible for tree pipit across-year and chiffchaff within-year. In contrast, larger positive differences were observed for tree pipit within-year (+1.6), chiffchaff across-year (+1.9), and great spotted kiwi (+2.6). The kiwi result showed the largest numerical difference, but it was not statistically significant, likely reflecting the higher inter-seed variability observed in the shuffled condition. These results suggest that chronological order may provide additional information in some longer or more structured vocalizations, but that this effect is not uniform across species or subset definitions.

### 3.5. Benchmarking with state-of-the-art

Table 6 summarizes performance relative to published results reported on the same datasets and compatible subset definitions. Details on protocol matching and metric alignment are provided in subsection 2.5. Values marked with ^†^ involve differences in split protocol and should therefore be interpreted as contextual benchmarks rather than direct like-for-like comparisons. In particular, the benchmarks based on the Stowell datasets used day-held-out or year-held-out evaluations, whereas our results are based on repeated stratified vocalization-level splits. All comparisons without markers correspond to protocol-compatible evaluations under comparable subset definitions.

**Table 6:** Comparison to published performance on the same datasets using compatible subset definitions. Results are grouped into classification and retrieval settings. Published values are reported as originally presented, whereas LSTM values are reported as mean ± standard deviation across five random seeds. Boldface indicates the best value within each species subset and metric when protocols are sufficiently comparable. RTBC: red-tailed black cockatoo; WY: within-year; AY: across-year.

| Species / subset | Reference | Model (reported) | Metric | Published | LSTM (Ours) |
| --- | --- | --- | --- | --- | --- |
| Classification setting |  |  |  |  |  |
| Chiffchaff (WY) | (Stowell et al., 2019) | CNN+RF | ROC-AUC | 0.9300<sup>†</sup> | 0.9996 ± 0.0002 |
|  | (Huang et al., 2025a) | AemNet | ROC-AUC | 0.9996 | 0.9996 ± 0.0002 |
|  | (Huang et al., 2025a) | AemNet | Accuracy | **0.9768** | 0.9747 ± 0.0076 |
| Chiffchaff (AY) | (Stowell et al., 2019) | CNN+RF | ROC-AUC | 0.5900<sup>†</sup> | 0.9998 ± 0.0001 |
| Little owl (AY) | (Stowell et al., 2019) | CNN+RF | ROC-AUC | 0.8500<sup>†</sup> | 0.9998 ± 0.0003 |
|  | (Huang et al., 2025a) | AemNet | ROC-AUC | 0.9961 | 0.9998 ± 0.0003 |
|  | (Huang et al., 2025a) | AemNet | Accuracy | 0.9204 | 0.9832 ± 0.0119 |
| Tree pipit (WY) | (Stowell et al., 2019) | CNN+RF | ROC-AUC | 0.9100<sup>†</sup> | 0.9995 ± 0.0008 |
|  | (Huang et al., 2025a) | AemNet | ROC-AUC | 0.9980 | 0.9995 ± 0.0008 |
|  | (Huang et al., 2025a) | AemNet | Accuracy | 0.9480 | 0.9706 ± 0.0239 |
| Tree pipit (AY) | (Stowell et al., 2019) | CNN+RF | ROC-AUC | 0.8300<sup>†</sup> | 0.9969 ± 0.0029 |
| Little penguin (WY) | (Huang et al., 2025a) | AemNet | ROC-AUC | 0.9976 | 0.9986 ± 0.0006 |
|  | (Huang et al., 2025a) | AemNet | Accuracy | 0.9363 | 0.9461 ± 0.0084 |
| RTBC (AY) | (Huang et al., 2025a) | AemNet | ROC-AUC | 0.9991 | 0.9992 ± 0.0006 |
|  | (Huang et al., 2025a) | AemNet | Accuracy | 0.9541 | 0.9659 ± 0.0048 |
| Great spotted kiwi (AY) | (Bedoya and Molles, 2021a) | KiwiNet+LAMDA | Accuracy (female) | 0.9460* | 0.9385 ± 0.0167 |
|  | (Bedoya and Molles, 2021a) | KiwiNet+LAMDA | Accuracy (male) | 0.8940* |  |
| Retrieval setting |  |  |  |  |  |
| Great tit (AY) | (Merino Recalde et al., 2025) | ViT emb+retrieval | CMC@1 | **0.9800** | 0.9689 ± 0.0018 |
|  | (Merino Recalde et al., 2025) | ViT emb+retrieval | mAP@5 | **0.9800** | 0.9800 ± 0.0011 |

Notes: <sup>†</sup> Protocol mismatch: published results used day-held-out or year-held-out splits, whereas our results use repeated stratified vocalization-level splits. \* Published values are sex-stratified with an unweighted mean of 92.0%; our results are a pooled closed-set accuracy on a balanced 20-individual set (10 females and 10 males).

Across the classification benchmarks, our framework (BirdNET + LSTM) achieved competitive performance relative to published baselines, although the strength of the comparison depends on protocol compatibility. The largest numerical differences occur in the Stowell-derived across-year subsets, where our ROC-AUC values are substantially higher than those reported for the CNN+RF pipeline. However, because these rows involve different split protocols, these differences should not be interpreted as direct evidence of superior across-year re-identification under the original chronological evaluation design. Relative to AemNet, performance remains broadly competitive. Our framework achieved equal or slightly higher ROC-AUC values across the matched comparisons, and higher accuracy for the little owl, tree pipit, little penguin, and red-tailed black cockatoo subsets, while AemNet retained a marginally higher accuracy for chiffchaff within-year.

For the great spotted kiwi, our experiment used a single pooled closed-set classification task including both sexes, with a balanced individual-level composition of 10 females and 10 males. However, although the kiwi subset was balanced at the individual level, the seed-specific test partitions were not sex-balanced at the vocalization level. Across the five seeds, female vocalizations represented 25–26 of 91 test samples, whereas male vocalizations represented 65–66 of 91 test samples. In contrast, Bedoya and Molles (2021a) reported sex-stratified accuracies separately for females (94.6%) and males (89.4%). As a contextual summary, the unweighted mean of these two sex-stratified accuracies is 92.0%, whereas our pooled 20-individual model achieved 93.9 ± 1.7%. Thus, our pooled accuracy should be interpreted as an overall closed-set estimate for the evaluated vocalization distribution, rather than as a sex-stratified estimate.

For the great tit, we report CMC@1 and mAP@5 computed from classifier scores as described in subsection 2.5. Under this evaluation, our framework achieves a comparable mAP@5 point estimate (0.9800 ± 0.0011 vs. 0.9800) with a decrease in CMC@1 (0.9689 ± 0.0018 vs. 0.9800), indicating similar top-*k* ranking quality despite different modeling paradigms (Merino Recalde et al., 2025).

## 4. DISCUSSION

### 4.1. Summary of contributions and main findings

This study shows that recurrent sequence modeling of bioacoustic embeddings can improve AIID in bird species whose vocalizations contain multiple informative BirdNET windows. By integrating transfer-learned BirdNET representations with a lightweight LSTM architecture, we demonstrate that individual acoustic signatures can be supported by acoustic information distributed across successive embedding windows. Across seven species and nine experimental conditions, the framework achieved mean test accuracies ranging from 93.9±1.7% to 98.3±0.4% and macro-F1 scores ranging from 93.2±0.9% to 98.3±1.3% across five stratified random seeds, without data augmentation. These results indicate that recurrent aggregation can capture identity-relevant information beyond simple static summaries in multi-window vocalizations, although the ordered-versus-shuffled ablation supports a cautious interpretation of temporal-order effects. To our knowledge, this is the first bioacoustic AIID study to combine pretrained foundation-model embeddings with recurrent sequence modeling at the individual level.

For deployment scenarios involving confidence thresholds or rejection rules, additional evaluation of score calibration may be warranted. This is relevant because macro-F1 and accuracy summarize closed-set argmax decisions, whereas ROC-AUC reflects score separation across thresholds. These metrics did not always vary in parallel, although numerical differences among configurations were generally small relative to the between-seed variability (see Supporting Information S2).

### 4.2. Sequence modeling, temporal order, and vocalization duration

The ablation results (Table 3) show that the benefit of recurrent sequence modeling was not uniform across species, but tended to be larger when vocalizations provided multiple informative BirdNET windows. The Onset baseline evaluated whether the first 3 s embedding was sufficient. In contrast, the comparison between the LSTM and GAP assessed whether recurrent nonlinear aggregation across all available embeddings improved performance beyond simple averaging. In this context, LSTM gains indicate that adaptive recurrent aggregation can improve identification in some cases, but they do not by themselves establish that chronological order is the driving factor.

This distinction is clearest in the longer-vocalization datasets. For the great spotted kiwi (27.64±3.92 s), the large gap between the Onset and GAP baselines shows that identity-relevant information was distributed beyond the first embedding. The additional improvement of the LSTM over GAP indicates that recurrent aggregation further improved performance over simple averaging. However, the ordered-versus-shuffled ablation (Table 5) showed that this improvement cannot be attributed uniformly to chronological order across all species. Preserving the original order produced a statistically significant gain only in the chiffchaff across-year subset, while the great spotted kiwi showed the largest mean ordered-shuffled difference, but with high inter-seed variability and no statistically significant effect. Thus, our results support the interpretation that multiple-window aggregation is important for some longer or more structured vocalizations, whereas the specific contribution of temporal order varied across the evaluated species and vocalization types.

For species whose vocalizations are shorter than, or close to, the 3 s BirdNET window, the proposed recurrent architecture offered limited or no demonstrated advantage over static baselines. In the red-tailed black cockatoo and little owl, vocalizations typically produced a single embedding, so the Onset and GAP baselines operated on the same input and were expected to converge. The LSTM also processed a one-step or near-one-step sequence in these cases, leaving little temporal structure to model. The little penguin showed a similar pattern, with the LSTM performing marginally below the best static baseline. Thus, these short-call datasets indicate no clear benefit or observed systematic degradation of recurrent modeling under single- or near-single-window conditions. From a practical perspective, a static classifier on top of BirdNET embeddings may be sufficient for high-accuracy identification when vocalizations are shorter than the embedding window, unless the segmentation or windowing strategy yields multiple informative embeddings.

The great tit represents a boundary case rather than a strictly short-call dataset. Although its mean duration was close to the 3 s BirdNET window, approximately one-third of test vocalizations generated more than one embedding. This limited amount of sequence-level information may explain why the LSTM showed a small but statistically significant gain over the strongest static baseline. However, the ordered-versus-shuffled comparison showed only a small non-significant effect for this subset, suggesting that the gain is more consistent with improved multiple-window aggregation than with strong evidence for temporal-order effects.

### 4.3. Pretrained representation choice and downstream aggregation

Our comparison of the BirdNET and Perch backbone and downstream aggregation configurations, summarized in Table 4, highlights a practical performance contrast among the evaluated AIID architectures. Under these evaluated configurations, the BirdNET + LSTM framework outperformed the Perch-based static baselines across all species and experimental conditions. The differences were small for the shortest vocalizations, for which both representations largely operated on a single embedding per vocalization. Larger differences emerged for the great tit, tree pipit, chiffchaff, and great spotted kiwi, suggesting that the combination of BirdNET embeddings with recurrent aggregation provided a stronger transfer-learning configuration for the AIID task on these datasets than Perch embeddings coupled with static downstream classifiers.

For vocalizations shorter than the Perch input window, such as the red-tailed black cockatoo, little owl, and little penguin, the Onset and GAP Perch baselines produced nearly identical results because each vocalization was typically represented by a single embedding. For intermediate-duration vocalizations, such as those of the great tit and tree pipit, the relative performance of Perch Onset and GAP varied across cases, suggesting that averaging across multiple windows was not consistently beneficial. One possible explanation is that additional windows may include a higher proportion of non-vocal or low-information audio when the vocalization occupies only part of the analysis window. However, because we did not explicitly quantify the proportion of padding or background content within each Perch window, this explanation should be treated as a plausible interpretation rather than as a demonstrated mechanism.

Finally, for species with vocalizations significantly longer than both window sizes, such as the great spotted kiwi (27.64 ± 3.92 s), the BirdNET + LSTM framework reached 95.1 ± 1.2% macro-F1, whereas the best Perch static baseline reached 91.7 ± 3.9%. This result indicates that, in this long-vocalization case, the evaluated BirdNET + LSTM configuration provided higher and more stable performance than the Perch-based static summaries. Nevertheless, because the comparison confounds the pretrained representation and the downstream aggregation strategy, it should be interpreted only as evidence of the practical effectiveness of the proposed configuration. A more direct separation of these factors would require applying recurrent classifiers to additional pretrained embedding backbones, including a Perch–LSTM condition, so that representation quality, input-window duration, and downstream sequence aggregation can be evaluated independently.

### 4.4. Generalization and robustness

A critical challenge in AIID is maintaining performance across temporal and ecological variation, since shifts in environmental conditions and animal behavior typically degrade model performance. Under our repeated stratified vocalization-level splits, the framework maintained high performance for species with both within-year and across-year subset definitions. For example, chiffchaff reached macro-F1 scores of 96.7 ± 1.0% in the within-year subset and 98.2 ± 0.4% in the across-year subset, while tree pipit reached 96.9±2.6% and 95.2±3.4%, respectively. However, these values should not be interpreted as strict evidence of temporal generalization to unseen years, because our splits were not year-held-out or session-disjoint.

This distinction is important when comparing with chronological benchmarks. For example, Stowell et al. (2019) reported a substantial decrease for chiffchaff when moving from within-season to across-season evaluation under a chronological split (ROC-AUC decreasing from 0.93 to 0.59). Our across-year results should therefore be interpreted as contextual benchmarks under repeated stratified partitioning, rather than as evidence that the framework resolves the stricter year-held-out re-identification problem. Nevertheless, they indicate that when seasonal variability is represented in the training data, the combination of BirdNET embeddings and LSTM-based aggregation can support individual identification of unseen vocalizations from known individuals.

Furthermore, our results remained competitive with published AIID approaches reported in Table 6, despite not using data augmentation. While previous studies often rely on noise injection, pitch shifting, or time-stretching to prevent overfitting to the training data (Stowell et al., 2019; Bedoya and Molles, 2021a; Merino Recalde et al., 2024), our framework achieved competitive performance in several matched or contextual comparisons using only transfer-learned embeddings. This suggests that substantial discriminative information is already present in the pretrained feature space and can become more separable when information distributed across multiple embedding windows is aggregated by a recurrent classifier, rather than requiring synthetic data expansion. These observations are consistent with Ghani et al. (2023), who showed that embeddings learned from large-scale bird-vocalization models support strong transfer learning in low-data regimes. Here, we extend that perspective from cross-class bioacoustic classification to within-species individual discrimination, showing that birdsong embeddings combined with recurrent aggregation can support strong performance on unseen vocalizations of known individuals under repeated stratified partitioning.

The framework also scaled well with population size. For the great tit, which had the largest class count (*N*_*c*_ = 242), the model achieved a macro-F1 of 94.0 ± 0.5% on the test set, despite the large number of individual classes. In the retrieval setting, our approach remained competitive with specialized architectures such as the ViT-based retrieval framework used by Merino Recalde et al. (2025). Although their model achieved higher top-1 accuracy (CMC@1: 0.9800 vs. our 0.9689 ± 0.0018), our LSTM-based framework matched it in mean average precision (mAP@5: our 0.9800 ± 0.0011 vs. 0.9800), indicating similarly strong ranking quality. Overall, these results show that the framework can reach competitive performance relative to state-of-the-art approaches without fine-tuning large-scale models or relying on complex data augmentation.

### 4.5. Limitations and threats to validity

While our results demonstrate the utility of coupling pretrained foundation-model embeddings with recurrent architectures for AIID, several limitations must be acknowledged regarding error propagation in the data pipeline, model interpretability, and ecological generalization.

First, the proposed framework is not an end-to-end detection-and-identification system; it relies on presegmented vocalizations. Consequently, performance is inherently bounded by the quality of the upstream segmentation process, as segmentation errors can truncate notes or alter temporal boundaries. This dependency was particularly critical for the great spotted kiwi dataset, where initial automated segmentation required refinement to ensure that the input sequences accurately captured the vocalization structure. In automated monitoring scenarios, poor segmentation could introduce temporal jitter or cut distinct notes, potentially degrading the sequential information available to the classifier.

Second, padding can dilute identity cues for short calls by reducing the effective signal content per window. Although the model showed robustness without data augmentation, the fixed input window strategy (3 s) presents a vulnerability for short signals. For short-call species, zero-padding increases the proportion of non-vocal content within the BirdNET window, potentially reducing the amount of active acoustic signal represented in the embedding. This suggests that even when recurrent aggregation can exploit information distributed across multiple windows, the underlying feature extractor (BirdNET) requires active signal presence to generate discriminative embeddings, and performance may suffer when short vocalizations contain substantial non-vocal background within the fixed analysis window. A related limitation is that non-overlapping 3 s windows may split syllables or notes at window boundaries, particularly when note timing is not aligned with the fixed BirdNET segmentation. Future work could evaluate overlapping 3 s windows with shorter strides to improve temporal resolution and reduce boundary effects, although this would increase computational cost and introduce stronger redundancy among adjacent embeddings.

Regarding label uncertainty, the little penguin dataset posed a unique challenge, as ground truth labels corresponded to nesting sites rather than biologically verified individuals. Despite this potential label noise (where a “nest” class might essentially represent a breeding pair), the system achieved a mean macro-F1 of 93.2 ± 0.9%. While this was the lowest score among the tested species, likely reflecting the inherent ambiguity of the labels and the limited amount of active vocal signal available within fixed-length windows, it nevertheless suggests the potential utility of the framework for monitoring stable social units or territories when individual tagging is impossible.

From a modeling perspective, a threat to validity arises from the domain mismatch of the backbone. BirdNET was optimized for species classification, a task that prioritizes inter-specific variance while potentially suppressing intra-specific (individual) variance to ensure generalization. While our results indicate that BirdNET embeddings retain sufficient individual information, it remains unknown whether these features are optimal. Furthermore, the use of LSTMs introduces a trade-off in interpretability. Unlike handcrafted feature analysis, where specific acoustic parameters (e.g., note duration, inter-syllable intervals) can be pinpointed as discriminative, the recurrent architecture operates as a “black box.” The individual-specific cues captured by the model are therefore not necessarily visible as simple differences in representative spectrograms, and may instead involve subtle combinations of frequency modulation, harmonic structure, syllable timing, and broader spectrotemporal patterns. Currently, we cannot quantify which temporal components, such as rhythm, phrase structure, or motif repetition, contribute to the identification decision.

A specific constraint regarding data independence must also be noted. Our primary evaluation used repeated stratified splits at the vocalization level rather than strictly session-disjoint partitions. Although this design allowed us to quantify sensitivity to random partitioning across five seeds, it does not ensure independence among recording sessions or years. Consequently, the reported performance should be interpreted as generalization to unseen vocalizations of known individuals under repeated stratified partitioning, and may remain optimistic relative to evaluations that explicitly test re-identification across independent sessions or years. This limitation is particularly relevant for comparisons with benchmarks based on chronological or session-held-out protocols.

The illustrative error analysis of the great spotted kiwi subset further showed that high aggregate accuracy can mask recurrent errors in a small number of individuals, highlighting the importance of inspecting class-level recall and confusion structure when adapting the workflow to deployment scenarios.

Finally, regarding ecological validity, our evaluation was conducted under a closed-set assumption, where all individuals in the test set were present during training. Real-world deployment, however, requires an open-set protocol capable of rejecting unknown individuals (“strangers”). Additionally, our datasets represent specific populations; generalization to other populations of the same species, which may exhibit local dialects, remains untested. Addressing these open-set and geographic generalization challenges is the necessary next step for deploying this framework in autonomous acoustic monitoring programs.

### 4.6. Practical implications and future work

The proposed methodology has practical implications for wildlife conservation and ethological monitoring by enabling non-invasive acoustic individual identification. In settings where capture and tagging are costly or undesirable, such as for threatened or elusive species like the great spotted kiwi, our framework provides an alternative route to assigning vocalizations to individuals from passive recordings. When paired with Autonomous Recording Units (ARUs), this approach can support long-term monitoring of individually identifiable callers while minimizing disturbance. In deployment, matching the analysis window and model complexity to the vocalization time scale can help avoid unnecessary computation and reduce windowing artifacts.

The computational efficiency of the proposed architecture further lowers barriers to adoption. All experiments were conducted on standard CPUs (4 vCPUs) without GPU acceleration, indicating that high-performing AIID models can be trained and evaluated on modest hardware. This reduces infrastructure requirements and makes deployment more feasible for conservation groups, NGOs, and research teams operating under limited computational budgets, particularly in resource-constrained regions.

To transition from a research pipeline to an autonomous field system, future work should address open-set identification. Realistic deployments require recognizing known individuals while rejecting previously unseen callers (“strangers”) and transient visitors. An intuitive next step is to integrate distance-based rejection or metric-learning objectives and to evaluate performance under explicit open-set protocols with calibrated thresholds. In parallel, integrating upstream Acoustic Event Detection (AED) would remove reliance on manual or semi-automatic segmentation, allowing raw ARU audio to be processed end-to-end into individual identity logs.

Finally, future work should improve the interpretability of temporal modeling. Attention-based pooling or Transformer-style sequence models could provide attribution over time and help identify which temporal cues (e.g., inter-note timing, phrase structure, or repetition patterns) contribute to individual discrimination. Optimizing the pipeline for edge computing may also enable on-device processing or near-real-time screening, reducing bandwidth demands and enabling timely identification alerts in the field.

## 5. CONCLUSIONS

Fixed-window approaches that summarize each vocalization into a single representation can constrain AIID for species with complex or extended vocalizations because they may fail to use information dis-tributed across multiple acoustic windows. We addressed this limitation by combining transfer-learned BirdNET embeddings with a lightweight LSTM that models sequences of embeddings. Our results show that individual identity can be encoded not only in short-window spectrotemporal content, but also in information distributed across successive embedding windows. However, the ablation experiment comparing ordered and shuffled embedding sequences indicates that the specific contribution of chronological order varied across the evaluated species and vocalization types, and should therefore be interpreted cautiously. Across seven species and nine experimental subsets from publicly available datasets used in prior AIID studies, the recurrent model achieved strong overall performance and improved over the strongest static baseline in the clearest long- or multi-window cases, particularly the great tit and great spotted kiwi subsets. Across five random seeds, test accuracy ranged from 93.9% to 98.3%, and macro-F1 ranged from 93.2% to 98.3%, without data augmentation. This indicates that discriminative identity cues are already present in the pretrained feature space and can become more separable when multiple embedding windows are aggregated with a recurrent classifier.

To our knowledge, this is the first study to combine bioacoustic foundation-model embeddings with recurrent sequence modeling for individual-level bird identification, yielding a CPU-friendly pipeline suitable for scalable monitoring workflows. The framework achieves competitive performance relative to published AIID approaches, relying only on pretrained embeddings and lightweight recurrent modeling, without requiring model fine-tuning or data augmentation. Our results also show that recurrent aggregation is most useful when vocalizations contain multiple informative windows, whereas for short vocalizations, the recurrent approach showed limited or no demonstrated advantage over simpler fully connected baselines. The comparison between BirdNET and Perch further suggests that pretrained representation choice and downstream aggregation strategy both affect performance, although these effects cannot be attributed solely to embedding-window duration. Finally, we highlight that workflow performance can be affected by upstream segmentation quality and by windowing choices that introduce low-information padded regions in short calls, which may dilute identity cues.

## AUTHOR CONTRIBUTIONS

**Jonathan Gallego:** Conceptualization, Methodology, Software, Validation, Formal analysis, Data curation, Writing – original draft, Visualization; **Juan D. Martínez:** Conceptualization, Software, Resources, Writing – review & editing; **José D. López:** Conceptualization, Validation, Writing – review & editing, Supervision, Project administration.

## DATA AVAILABILITY STATEMENT

All materials required to reproduce the analyses are archived in a permanent repository. The complete data package (datasets repackaged into the directory structure required by the code, metadata including split files, the kiwi trimming notebook, and precomputed embeddings in Parquet format for BirdNET and Perch) is available on Zenodo: https://doi.org/10.5281/zenodo.18603175. Source code is hosted across two repositories: preprocessing, including temporal normalization (padding), and embedding extraction using birdnetlib (https://github.com/jongalon/emb_extraction) and model training/evaluation (LSTM and fully connected baselines, plus ablation logic) (https://github.com/jongalon/embedding-to-individual-id).

## ACKNOWLEDGMENTS

This research was supported by Universidad de Antioquia – CODI and Rice University (code project: 2024-73410)

## SUPPORTING INFORMATION

Additional supporting information can be found online in the Supporting Information section at the end of this article. **Supporting Information S1:** Additional segmentation algorithm for great spotted kiwi. **Supporting Information S2:** Additional performance metrics for all baseline configurations. **Supporting Information S3:** Illustrative per-individual error analysis for the great spotted kiwi subset.

