## Supporting Information S1 for "Individual Bird Identification by Modeling Temporal Structure in Bioacoustic Embeddings"

### Supporting Information S1: Kiwi vocalization trimming procedure

This document describes the trimming procedure used to reduce background audio around kiwi vocalizations in the great spotted kiwi dataset, where source clips were provided as 60 s segments. The goal is to produce shorter clips centered on the detected vocal activity by removing leading and trailing low-energy regions, while preserving a small padding margin around vocalizations.

#### S1.1 Inputs and outputs

Input: WAV audio files (mono or multichannel) stored under an input directory. If an input file has multiple channels, channels are averaged to mono prior to detection.

Output: trimmed WAV files saved under an output directory while preserving the input subfolder structure. In the default configuration (export_mode = trim_edges), each file produces one output named <stem>_trim.wav. A global CSV log (trimming_log.csv) is written to the output directory summarizing the trimming bounds and number of detected segments per file.

#### S1.2 Parameter configuration

Table S1 lists the parameters used in the batch run that generated the trimmed kiwi clips.

| Parameter | Value | Description |
|---|---|---|
| method | "energy" | Detection backend used for trimming. |
| export_mode | "trim_edges" | Output mode: trim to a single continuous interval or export each detected segment. |
| target_sr | None | Optional resampling target sample rate. None keeps the original sampling rate. |
| ANALYSIS_ROI | (5.0, 55.0) | Time interval (s) used for detection only. The final trimming is applied on |

| | | |
|---|---|---|
| | | the full recording using the detected bounds. |
| bandpass_low_hz | 1000 | Lower cutoff frequency (Hz) for Butterworth band-pass filtering before RMS computation. |
| bandpass_high_hz | 3600 | Upper cutoff frequency (Hz) for Butterworth band-pass filtering before RMS computation. |
| frame_length | 1024 | Frame length (samples) for RMS computation. |
| hop_length | 256 | Hop length (samples) for RMS computation. |
| smooth_win_sec | 0.20 | Median-smoothing window (seconds) applied to RMS for robust noise-floor estimation. |
| noise_strategy | "quantile" | Noise-set construction strategy. quantile uses the lowest-energy portion of RMS as noise. |
| noise_q | 0.50 | Quantile used to select low-energy frames for the noise set when noise_strategy is quantile. |
| threshold_mode | "percentile" | Thresholding rule. percentile uses a percentile of the noise-set RMS. |
| noise_percentile | 98.0 | Percentile of the noise-set RMS used as threshold when threshold_mode is percentile. |

| | | |
|---|---|---|
| k_std | 5.0 | STD multiplier used when threshold_mode is std. |
| min_segment_sec | 0.5 | Minimum segment duration (s) kept after thresholding. |
| merge_gap_sec | 0.1 | Silent gap (s) below which adjacent segments are merged. |
| prepad_sec | 0.10 | Padding (s) added before the first detected segment (or each segment in all_segments mode). |
| postpad_sec | 0.10 | Padding (s) added after the last detected segment (or each segment in all_segments mode). |
| ENABLE_CLUSTER_FILTER | True | If enabled, keeps only the main cluster of segments and suppresses stray detections. |
| BIG_GAP_SEC | 4.5 | Temporal gap (s) that defines a new cluster of segments. |
| MAIN_CLUSTER_MODE | "duration" | Criterion to select the main cluster (duration, span, or count). |
| MIN_MAIN_CLUSTER_NOTES | 6 | Guard: minimum number of segments required for the chosen main cluster. |
| MIN_MAIN_CLUSTER_DURATION | 5.0 | Guard: minimum total duration (s) required for the chosen main cluster. |
| DROP_ISOLATED_TINY | True | If enabled, drops isolated very short segments before clustering. |

| | | |
|---|---|---|
| ISOLATION_GAP_SEC | 2.0 | Isolation gap (s) used to mark a segment as isolated. |
| MAX_TINY_SEC | 0.7 | Maximum duration (s) for a segment to be considered a tiny isolated blip. |
| BATCH_QC_EVERY | 1 | QC plot frequency. 1 saves a QC plot for every file. |

### S1.3 Trimming algorithm

The batch procedure proceeds as follows:

1. Load audio using soundfile, convert to mono by channel averaging if needed, and cast to float32.
2. Define an analysis region of interest (ROI) in seconds (ANALYSIS_ROI). Detection is performed only on this ROI, but trimming is applied on the full recording based on the detected bounds.
3. Optionally band-pass filter the ROI using a 4th-order Butterworth filter and zero-phase forward-backward filtering (filtfilt).
4. Compute a frame-wise RMS envelope using frame_length and hop_length.
5. Apply median smoothing to the RMS envelope over smooth_win_sec to stabilize the noise-floor estimate.
6. Construct a noise set using the lowest-energy portion of frames (noise_strategy = quantile, noise_q).
7. Compute an RMS threshold. With threshold_mode = percentile, the threshold is the noise_percentile of the noise-set RMS distribution. Frames with RMS greater than the threshold are marked as voiced.
8. Convert the voiced mask into contiguous voiced segments. Merge gaps shorter than merge_gap_sec and drop segments shorter than min_segment_sec.
9. Shift detected segments from ROI time to global time (add roi_start). If enabled, suppress stray detections by clustering segments separated by BIG_GAP_SEC and retaining only the main cluster (selected by MAIN_CLUSTER_MODE). Optionally drop isolated tiny segments before clustering (DROP_ISOLATED_TINY, ISOLATION_GAP_SEC, MAX_TINY_SEC).
10. Fallback: if no segments are detected, retry percentile thresholding with progressively lower noise_percentile values (subtracting 3, 6, and 9) until at least one segment is detected or a lower bound is reached.
11. Export trimmed audio. For trim_edges, compute bounds from the first and last detected segment and add prepad_sec and postpad_sec (clipped to the recording duration), then

write one trimmed file. For all_segments, export each detected segment separately with padding.

12. Optionally produce QC plots (RMS, threshold, ROI, detected segments, and final bounds) at the configured frequency (BATCH_QC_EVERY) and save them under the QC directory.

#### S1.4 Pseudocode

```
for each file in input_dir:
    y, sr = read_audio(file)
    y = to_mono(y)
    y = float32(y)

    roi_start, roi_end = clamp(ANALYSIS_ROI, 0, duration(y, sr))
    y_roi = y[roi_start:roi_end]

    segments = detect_voiced_regions(
        y_roi, sr,
        frame_length, hop_length,
        bandpass_low_hz, bandpass_high_hz,
        noise_strategy, noise_q,
        threshold_mode, noise_percentile,
        min_segment_sec, merge_gap_sec,
        smooth_win_sec
    )
    segments = shift_to_global_time(segments, roi_start)

    if ENABLE_CLUSTER_FILTER:
        segments = post_cluster_filter(segments, BIG_GAP_SEC,
MAIN_CLUSTER_MODE,
                                       MIN_MAIN_CLUSTER_NOTES,
MIN_MAIN_CLUSTER_DURATION,
                                       DROP_ISOLATED_TINY, ISOLATION_GAP_SEC,
MAX_TINY_SEC)

    if segments is empty and threshold_mode == "percentile":
        retry with noise_percentile - {3, 6, 9}

    if segments is empty:
        log "none_found" and continue

    if export_mode == "trim_edges":
        bounds = [first_start - prepad_sec, last_end + postpad_sec]
        write y[bounds] to <stem>_trim.wav
    else:
        write each segment with padding to <stem>_segXX.wav

    append record to trimming_log.csv
```

### S1.5 Worked example: ROI selection and trimming outcome

Figures S1 and S2 provide representative outputs from the trimming procedure. Figure S1 shows the ROI-based RMS envelope, detection threshold, and derived trimming bounds for one 60 s clip. Figure S2 compares the spectrogram before and after trimming for the same recording.

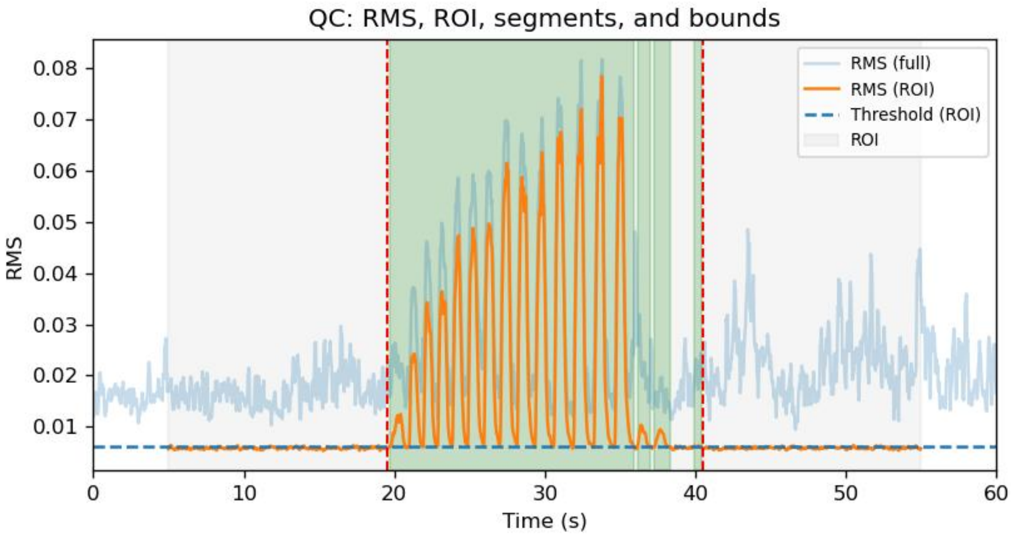


*Figure S1. Quality-control plot for one 60 s clip. The RMS envelope is computed within the ROI (gray shading), compared against the detection threshold (blue dashed line), and used to derive trimming bounds (red dashed lines).*

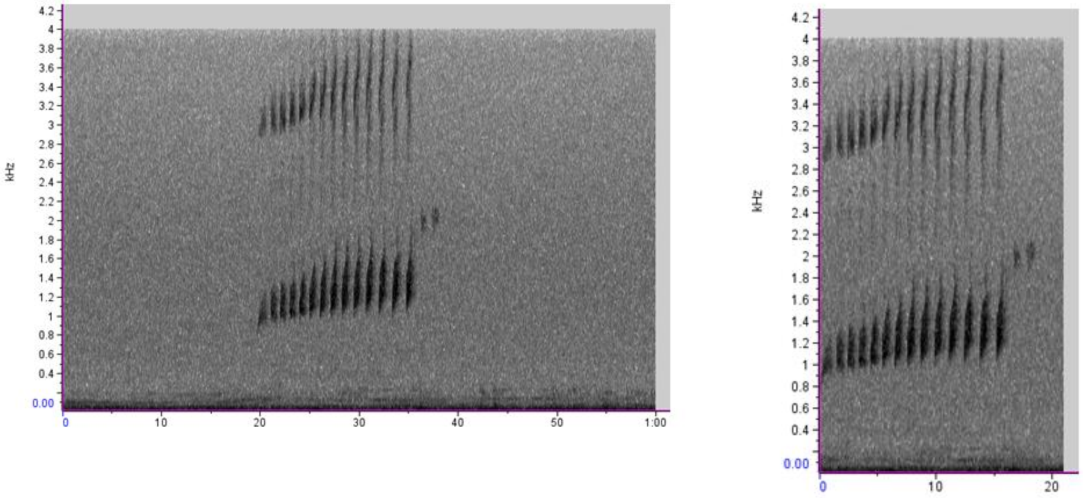


*Figure S2. Spectrogram comparison for the same recording. Left: original 60 s clip. Right: trimmed clip produced by the algorithm.*

#### S1.6 Notes on reproducibility

The procedure was implemented as a Jupyter notebook to facilitate iterative development and quality control through visual inspection. Access details are provided in the manuscript Data Availability section.
