## Supporting Information S2 for "Individual Bird Identification by Modeling Temporal Structure in Bioacoustic Embeddings"

### Supporting Information S2: Performance metrics for all baseline configurations.

This document provides a comprehensive summary of classification performance across all evaluated model configurations. For each species and experimental subset, we report macro-F1, accuracy, and ROC-AUC as mean ± standard deviation over five random seeds, to complement the main-text results and to facilitate direct comparison among approaches. Configurations include the BirdNET-based variants (static aggregation baselines and the recurrent BirdNET + LSTM model) as well as Perch-based baselines where applicable. The final table additionally compares the ordered and shuffled BirdNET + LSTM variants. Metrics are presented in a consistent tabular format and grouped by species and subset to highlight how performance varies with different data splits and vocalization conditions. These additional metrics are included to support transparency, enable reproducibility, and allow readers to assess whether observed differences are robust across complementary evaluation criteria.

**Table S2.1. BirdNET-based ablation results (BirdNET + Onset, BirdNET + GAP, BirdNET + LSTM). Metrics are reported as mean ± standard deviation over five random seeds. Macro-F1 and Accuracy are in percentages; ROC-AUC is reported on the [0,1] scale. Boldface indicates the best Macro-F1 across the three configurations within each species/subset.**

| Species | Dur. (s)/Voc | Subset | Model | Macro-F1 (%) | Accuracy (%) | ROC-AUC | Vocs/indiv |
|---|---|---|---|---|---|---|---|
| Chiffchaff (*P. collybita*) | 5.35 ± 2.56 | within-year | BirdNET + Onset | 96.0 ± 0.3 | 96.9 ± 0.2 | 0.9993 ± 0.0002 | 480 ± 272 |
| | | | BirdNET + GAP | 96.6 ± 1.4 | 97.4 ± 0.9 | 0.9996 ± 0.0002 | |
| | | | BirdNET + LSTM | **96.7 ± 1.0** | 97.5 ± 0.8 | 0.9996 ± 0.0002 | |
| | | across-year | BirdNET + Onset | 94.7 ± 2.5 | 94.5 ± 2.7 | 0.9988 ± 0.0006 | 52 ± 13 |
| | | | BirdNET + GAP | 96.6 ± 1.7 | 96.6 ± 1.7 | 0.9992 ± 0.0005 | |
| | | | BirdNET + LSTM | **98.2 ± 0.4** | 98.3 ± 0.4 | 0.9998 ± 0.0001 | |
| Little owl (*A. noctua*) | 1.23 ± 0.23 | across-year | BirdNET + Onset | 97.9 ± 1.0 | 97.9 ± 1.0 | 0.9997 ± 0.0003 | 60 ± 8 |
| | | | BirdNET + GAP | 97.9 ± 1.0 | 97.9 ± 1.0 | 0.9997 ± 0.0003 | |
| | | | BirdNET + LSTM | **98.3 ± 1.3** | 98.3 ± 1.2 | 0.9998 ± 0.0003 | |
| Tree pipit (*A. trivialis*) | 4.03 ± 2.09 | within-year | BirdNET + Onset | 94.0 ± 3.7 | 94.5 ± 3.0 | 0.9986 ± 0.0013 | 71 ± 16 |
| | | | BirdNET + GAP | 95.0 ± 3.1 | 95.4 ± 2.7 | 0.9982 ± 0.0015 | |
| | | | BirdNET + LSTM | **96.9 ± 2.6** | 97.1 ± 2.4 | 0.9995 ± 0.0008 | |
| | | across-year | BirdNET + Onset | **95.8 ± 2.0** | 95.7 ± 2.0 | 0.9986 ± 0.0011 | 72 ± 15 |
| | | | BirdNET + GAP | 95.5 ± 1.9 | 95.7 ± 1.6 | 0.9979 ± 0.0012 | |
| | | | BirdNET + LSTM | 95.2 ± 3.4 | 95.4 ± 3.1 | 0.9969 ± 0.0029 | |
| Little penguin (*E. minor*) | 1.95 ± 0.77 | within-year | BirdNET + Onset | **93.4 ± 0.9** | 94.2 ± 1.6 | 0.9983 ± 0.0008 | 162 ± 71 |
| | | | BirdNET + GAP | 93.1 ± 1.1 | 93.9 ± 0.7 | 0.9984 ± 0.0005 | |
| | | | BirdNET + LSTM | 93.2 ± 0.9 | 94.6 ± 0.8 | 0.9986 ± 0.0006 | |
| Red-tailed black cockatoo (*C. banksii*) | 0.76 ± 0.17 | across-year | BirdNET + Onset | 94.7 ± 3.7 | 96.4 ± 0.9 | 0.9991 ± 0.0007 | 112 ± 116 |
| | | | BirdNET + GAP | 94.7 ± 3.7 | 96.4 ± 0.9 | 0.9991 ± 0.0007 | |
| | | | BirdNET + LSTM | **95.9 ± 1.2** | 96.6 ± 0.5 | 0.9992 ± 0.0006 | |
| Great spotted kiwi (*A. maxima*) | 27.64 ± 3.92 | across-year | BirdNET + Onset | 79.0 ± 3.6 | 83.7 ± 2.9 | 0.9893 ± 0.0038 | 23 ± 24 |
| | | | BirdNET + GAP | 92.3 ± 0.8 | 92.3 ± 0.8 | 0.9987 ± 0.0008 | |
| | | | BirdNET + LSTM | **95.1 ± 1.2** | 93.8 ± 1.7 | 0.9982 ± 0.0028 | |
| Great tit (*P. major*) | 2.66 ± 1.13 | across-year | BirdNET + Onset | 92.6 ± 0.9 | 96.1 ± 0.5 | 0.9997 ± 0.0002 | 306 ± 314 |
| | | | BirdNET + GAP | 92.3 ± 1.1 | 95.7 ± 0.6 | 0.9996 ± 0.0002 | |
| | | | BirdNET + LSTM | **94.0 ± 0.5** | 96.9 ± 0.2 | 0.9998 ± 0.0001 | |

**Table S2.2. Backbone comparison including Perch-based baselines (Perch + Onset, Perch + GAP) and the recurrent BirdNET model (BirdNET + LSTM). Values correspond to the same species and subsets as Table S1 and follow the same reporting convention (mean ± standard deviation over five random seeds). Boldface indicates the best Macro-F1 across the three configurations within each species/subset.**

| Species | Dur. (s)/Voc | Subset | Model | Macro-F1 (%) | Accuracy (%) | ROC-AUC | Vocs/indiv |
|---|---|---|---|---|---|---|---|
| Chiffchaff (*P. collybita*) | 5.35 ± 2.56 | within-year | Perch + Onset | 92.3 ± 1.4 | 93.8 ± 1.1 | 0.9971 ± 0.0009 | |
| | | | Perch + GAP | 92.3 ± 0.6 | 93.8 ± 0.3 | 0.9973 ± 0.0005 | 480 ± 272 |
| | | | BirdNET + LSTM | **96.7 ± 1.0** | 97.5 ± 0.8 | 0.9996 ± 0.0002 | |
| | | across-year | Perch + Onset | 89.5 ± 3.3 | 89.0 ± 3.0 | 0.9926 ± 0.0034 | |
| | | | Perch + GAP | 83.3 ± 4.9 | 84.2 ± 4.3 | 0.9856 ± 0.0102 | 52 ± 13 |
| | | | BirdNET + LSTM | **98.2 ± 0.4** | 98.3 ± 0.4 | 0.9998 ± 0.0001 | |
| Little owl (*A. noctua*) | 1.23 ± 0.23 | across-year | Perch + Onset | 97.5 ± 0.3 | 97.4 ± 0.4 | 0.9995 ± 0.0003 | |
| | | | Perch + GAP | 97.5 ± 0.3 | 97.4 ± 0.4 | 0.9995 ± 0.0003 | 60 ± 8 |
| | | | BirdNET + LSTM | **98.3 ± 1.3** | 98.3 ± 1.2 | 0.9998 ± 0.0003 | |
| Tree pipit (*A. trivialis*) | 4.03 ± 2.09 | within-year | Perch + Onset | 89.8 ± 3.6 | 90.9 ± 3.1 | 0.9939 ± 0.0045 | |
| | | | Perch + GAP | 90.4 ± 3.4 | 91.5 ± 3.1 | 0.9924 ± 0.0036 | 71 ± 16 |
| | | | BirdNET + LSTM | **96.9 ± 2.6** | 97.1 ± 2.4 | 0.9995 ± 0.0008 | |
| | | across-year | Perch + Onset | 90.5 ± 2.3 | 91.0 ± 2.0 | 0.9926 ± 0.0060 | |
| | | | Perch + GAP | 89.9 ± 2.4 | 90.8 ± 2.1 | 0.9915 ± 0.0035 | 72 ± 15 |
| | | | BirdNET + LSTM | **95.2 ± 3.4** | 95.4 ± 3.1 | 0.9969 ± 0.0029 | |
| Little penguin (*E. minor*) | 1.95 ± 0.77 | within-year | Perch + Onset | 92.2 ± 1.0 | 92.6 ± 0.8 | 0.9971 ± 0.0008 | |
| | | | Perch + GAP | 92.2 ± 1.0 | 92.6 ± 0.8 | 0.9971 ± 0.0008 | 162 ± 71 |
| | | | BirdNET + LSTM | **93.2 ± 0.9** | 94.6 ± 0.8 | 0.9986 ± 0.0006 | |
| Red-tailed black cockatoo (*C. banksii*) | 0.76 ± 0.17 | across-year | Perch + Onset | 95.0 ± 2.2 | 96.2 ± 0.6 | 0.9993 ± 0.0005 | |
| | | | Perch + GAP | 95.0 ± 2.2 | 96.2 ± 0.6 | 0.9993 ± 0.0005 | 112 ± 116 |
| | | | BirdNET + LSTM | **95.9 ± 1.2** | 96.6 ± 0.5 | 0.9992 ± 0.0006 | |
| Great spotted kiwi (*A. maxima*) | 27.64 ± 3.92 | across-year | Perch + Onset | 84.8 ± 4.8 | 86.6 ± 2.4 | 0.9925 ± 0.0034 | |
| | | | Perch + GAP | 91.7 ± 3.9 | 92.5 ± 3.0 | 0.9989 ± 0.0009 | 23 ± 24 |
| | | | BirdNET + LSTM | **95.1 ± 1.2** | 93.8 ± 1.7 | 0.9982 ± 0.0028 | |
| Great tit (*P. major*) | 2.66 ± 1.13 | across-year | Perch + Onset | 85.9 ± 0.8 | 91.5 ± 0.6 | 0.9989 ± 0.0003 | |
| | | | Perch + GAP | 86.3 ± 0.9 | 91.5 ± 0.5 | 0.9988 ± 0.0003 | 306 ± 314 |
| | | | BirdNET + LSTM | **94.0 ± 0.5** | 96.9 ± 0.2 | 0.9998 ± 0.0001 | |

**Table S2.3. Effect of temporal ordering on the BirdNET + LSTM model. Comparison between the ordered LSTM (chronologically sorted embedding sequences) and the shuffled LSTM (seed-specific random permutations of the same sequences). Reported values are mean ± standard deviation over five random seeds. Only species with multiple embeddings per vocalization are included, since shuffling has no effect when each vocalization is represented by a single embedding (little owl, little penguin, red-tailed black cockatoo). Boldface indicates the higher Macro-F1 between ordered and shuffled within each species/subset.**

| **Species** | **Dur. (s)/Voc** | **Subset** | **Sequence** | **Macro-F1 (%)** | **Accuracy (%)** | **ROC-AUC** |
|---|---|---|---|---|---|---|
| Great tit (*P. major*) | 2.66 ± 1.13 | across-year | Ordered | **94.0 ± 0.5** | 96.9 ± 0.2 | 0.9998 ± 0.0001 |
| | | | Shuffled | 93.7 ± 0.5 | 96.7 ± 0.3 | 0.9998 ± 0.0001 |
| Tree pipit (*A. trivialis*) | 4.03 ± 2.09 | within-year | Ordered | **96.9 ± 2.6** | 97.1 ± 2.4 | 0.9995 ± 0.0008 |
| | | | Shuffled | 95.4 ± 2.6 | 95.8 ± 2.1 | 0.9987 ± 0.0008 |
| | | across-year | Ordered | **95.2 ± 3.4** | 95.4 ± 3.1 | 0.9969 ± 0.0029 |
| | | | Shuffled | 95.1 ± 1.3 | 95.3 ± 1.3 | 0.9975 ± 0.0015 |
| Chiffchaff (*P. collybita*) | 5.35 ± 2.56 | within-year | Ordered | 96.7 ± 1.0 | 97.5 ± 0.8 | 0.9996 ± 0.0002 |
| | | | Shuffled | **96.8 ± 0.4** | 97.4 ± 0.4 | 0.9995 ± 0.0002 |
| | | across-year | Ordered | **98.2 ± 0.4** | 98.3 ± 0.4 | 0.9998 ± 0.0001 |
| | | | Shuffled | 96.3 ± 1.2 | 96.2 ± 1.5 | 0.9994 ± 0.0003 |
| Great spotted kiwi (*A. maxima*) | 27.64 ± 3.92 | across-year | Ordered | **95.1 ± 1.2** | 93.8 ± 1.7 | 0.9982 ± 0.0028 |
| | | | Shuffled | 92.5 ± 3.8 | 91.6 ± 3.9 | 0.9980 ± 0.0022 |
