## Supporting Information S3 for "Individual Bird Identification by Modeling Temporal Structure in Bioacoustic Embeddings"

### Supporting Information S3: Illustrative per-individual error analysis for the great spotted kiwi subset.

This document provides an illustrative per-individual error analysis for the great spotted kiwi across-year subset, using the ordered BirdNET + LSTM model evaluated over five random seeds. The aim was not to develop a full error taxonomy across all species, but to examine whether the remaining misclassifications in a long-vocalization subset were concentrated in particular random partitions, individuals with limited training data, or recurrent true-predicted confusion pairs. We first summarize the total number of test predictions, errors, and accuracy for each seed (Table S3.1). We then aggregate errors at the individual level across seeds to examine whether failures were concentrated in specific individuals (Table S3.2). To assess whether these class-level errors were associated with the amount of available training data, we report Spearman correlations between mean training-set size and both pooled recall and error rate (Table S3.3). Finally, we list the most frequent recurrent confusion pairs to identify whether some errors reflected repeated misassignments between the same individuals rather than isolated random mistakes (Table S3.4).

For each seed, test-set predictions were matched with the corresponding seed-specific split assignments. We computed seed-level accuracy and error counts, pooled per-individual recall across seeds, the mean number of training vocalizations available for each individual, recurrent true-predicted confusion pairs, and Spearman correlations between training-set size and per-individual performance.

**Table S3.1. Seed-level error summary for the great spotted kiwi across-year subset using the ordered BirdNET + LSTM model. Test predictions, errors, and accuracy are reported separately for each random seed and pooled across the five seeds.**

| Seed | Test predictions | Errors | Accuracy (%) |
|---|---|---|---|
| 18 | 91 | 5 | 94.51 |
| 23 | 91 | 4 | 95.60 |
| 46 | 91 | 6 | 93.41 |
| 123 | 91 | 8 | 91.21 |
| 321 | 91 | 5 | 94.51 |
| **Total** | **455** | **28** | **93.85** |

At the seed level, accuracy ranged from 91.21% to 95.60%, with 28 errors across 455 pooled test predictions (Table S3.1). This indicates that performance was consistently high across partitions, although the number of residual errors varied moderately among seeds.

**Table S3.2. Per-individual error summary for the great spotted kiwi across-year subset aggregated across five random seeds. Mean train n denotes the average number of training vocalizations per individual across seed-specific partitions. Test n, errors, and pooled recall were computed from all test predictions across seeds.**

| Individual | Mean train n | Test n | Errors | Pooled recall |
|---|---|---|---|---|
| M9 | 68.0 | 115 | 8 | 0.930 |
| M7 | 20.0 | 30 | 5 | 0.833 |
| M99 | 32.0 | 55 | 4 | 0.927 |
| M5 | 9.0 | 15 | 2 | 0.867 |
| M4 | 12.0 | 20 | 2 | 0.900 |
| M8 | 22.0 | 35 | 2 | 0.943 |
| F7 | 6.0 | 10 | 1 | 0.900 |
| F9 | 5.0 | 10 | 1 | 0.900 |
| F5 | 7.8 | 12 | 1 | 0.917 |
| M6 | 7.8 | 14 | 1 | 0.929 |
| F6 | 10.8 | 20 | 1 | 0.950 |
| F1 | 10.0 | 15 | 0 | 1.000 |
| F2 | 8.0 | 15 | 0 | 1.000 |
| F3 | 6.0 | 10 | 0 | 1.000 |
| F4 | 10.0 | 15 | 0 | 1.000 |
| F8 | 4.8 | 8 | 0 | 1.000 |
| F99 | 8.0 | 13 | 0 | 1.000 |
| M1 | 7.8 | 13 | 0 | 1.000 |
| M2 | 13.0 | 20 | 0 | 1.000 |
| M3 | 5.0 | 10 | 0 | 1.000 |

The per-individual summary showed that the 28 errors were not uniformly distributed across individuals (Table S3.2). Eleven of the 20 individuals had at least one error, while the three individuals with the most errors accounted for 60.7% of all errors, and the five individuals with the most errors accounted for 75.0%. However, errors were not explained solely by training-set size. Some individuals with relatively large training sets, such as M9 and M99, accumulated multiple errors, whereas several individuals with smaller training sets had no errors across the five seeds.

**Table S3.3. Relationship between training-set size and per-individual performance in the great spotted kiwi error analysis. Spearman correlations were computed across individuals after aggregating predictions across the five random seeds.**

| Comparison | Spearman ρ | p-value |
|---|---|---|
| Mean train n vs. pooled recall | -0.211 | 0.373 |
| Mean train n vs. error rate | 0.211 | 0.373 |

The correlation analysis provided no evidence that individuals with fewer training vocalizations systematically had lower recall or higher error rates (Table S3.3). The weak and non-significant correlations suggest that the observed class-level errors were not primarily driven by training-set size, although the small number of individuals limits the strength of this inference.

**Table S3.4. Most frequent recurrent true-predicted confusion pairs in the great spotted kiwi across-year subset. Pairs are aggregated across the five random seeds, and only pairs with at least two errors are shown.**

| True individual | Predicted individual | Errors | Seeds |
|---|---|---|---|
| M9 | M8 | 5 | 46, 123, 321 |
| M9 | M99 | 3 | 23, 46 |
| M99 | M9 | 3 | 23, 123 |
| M7 | M8 | 2 | 18, 123 |
| M4 | M8 | 2 | 18 |
| M5 | M99 | 2 | 23, 123 |
| M7 | M99 | 2 | 46, 123 |
| M8 | M7 | 2 | 321 |

The recurrent confusion-pair analysis showed that several errors occurred repeatedly between the same individuals across seeds (Table S3.4). For example, confusions involving M9, M8, and M99 accounted for multiple repeated errors, suggesting that at least part of the residual error structure reflected systematic similarity among some individuals rather than purely random misclassification.

Overall, this illustrative analysis indicates that high aggregate accuracy can mask recurrent class-level errors. In the great spotted kiwi subset, residual misclassifications were not solely explained by the number of available training vocalizations per individual, but were partially concentrated in a small number of individuals and recurrent confusion pairs. These results support inspecting class-level recall and confusion structure when adapting the workflow to deployment-oriented monitoring settings.
